# Niche differentiation confers coexistence prior to the species boundary in an aquatic plant

**DOI:** 10.64898/2026.04.03.716288

**Authors:** Jawad Sakarchi, Takuji Usui, Pablo Duchen, Simon P. Hart, Martin M. Turcotte, Shuqing Xu, Amy L. Angert, Rachel M. Germain

## Abstract

Despite the prevailing view that ecological divergence drives speciation, we know little about when or how nascent lineages evolve the ecological differences needed to coexist upon secondary contact. Here, we apply ecological coexistence theory to quantify the potential for coexistence among 126 allopatric lineages of the globally distributed duckweed *Spirodela polyrhiza*. Using competition experiments simulating secondary contact, we found that nascent lineages were more likely to coexist as they become genetically distinct. This occurred through the rise of niche differences, but not competitive differences, early in lineage divergence. Competition assays against its sister-species *Spirodela intermedia* further showed that most niche differences may already be in place to promote coexistence well before timescales at which speciation is complete. Our findings suggest that early coexistence may thus create the ecological conditions for sympatry among nascent lineages and broaden opportunities for gene flow during speciation, serving as an important but overlooked link between the ecological controls that maintain biodiversity and its evolutionary origins.

## Main Text

The immense biodiversity we see on Earth today represents the rare successes of speciation out of many failures ^1–3^. Despite a long history of speciation research focusing on the evolution of reproductive isolation ^2^, there is growing evidence that reproductive isolation need not be the rate-limiting step ^4–10^. Recent studies highlight how the accumulation of species diversity could also hinge upon whether ecological differentiation, if any, could permit diverging lineages to demographically coexist as separate species ^1,11–13^ Under the dominant view that lineage divergence proceeds largely in allopatry ^14^, populations separated over greater evolutionary time and across more divergent environments are more likely to be both reproductively isolated and ecologically differentiated upon secondary contact ^14–16^. Yet, whether ecological differentiation acts to stabilize the coexistence of allopatric populations if they were to come into contact, and if mechanisms enabling lineage coexistence evolve at timescales relevant to affect speciation, is unknown.

This presents a key gap in our understanding of the ecological controls for why and when speciation could succeed or fail, as both the timescale at which coexistence evolves relative to reproductive isolation (i.e., tempo) and the mechanism by which lineages ecologically differentiate (i.e., mode) could alter the outcome of speciation (Fig. 1A). For tempo, the classic view is that lineages in allopatry evolve mechanisms to coexist after reproductive isolation is established, promoting the completion of speciation by allowing reproductively isolated lineages to persist even upon secondary contact ^2,3,11,14^ However, if lineages evolve the potential to coexist prior to the evolution of reproductive isolation, then hybridization may cause nascent species to collapse back into one, halting speciation ^3,11,17^ As ecological coexistence and its evolution is classically studied across distinct species at macroevolutionary timescales ^18–20^, empirical data on if and how fast ecological differences accumulate among nascent lineages to influence coexistence within species and across the speciation boundary is critically lacking.

**Fig. 1.**
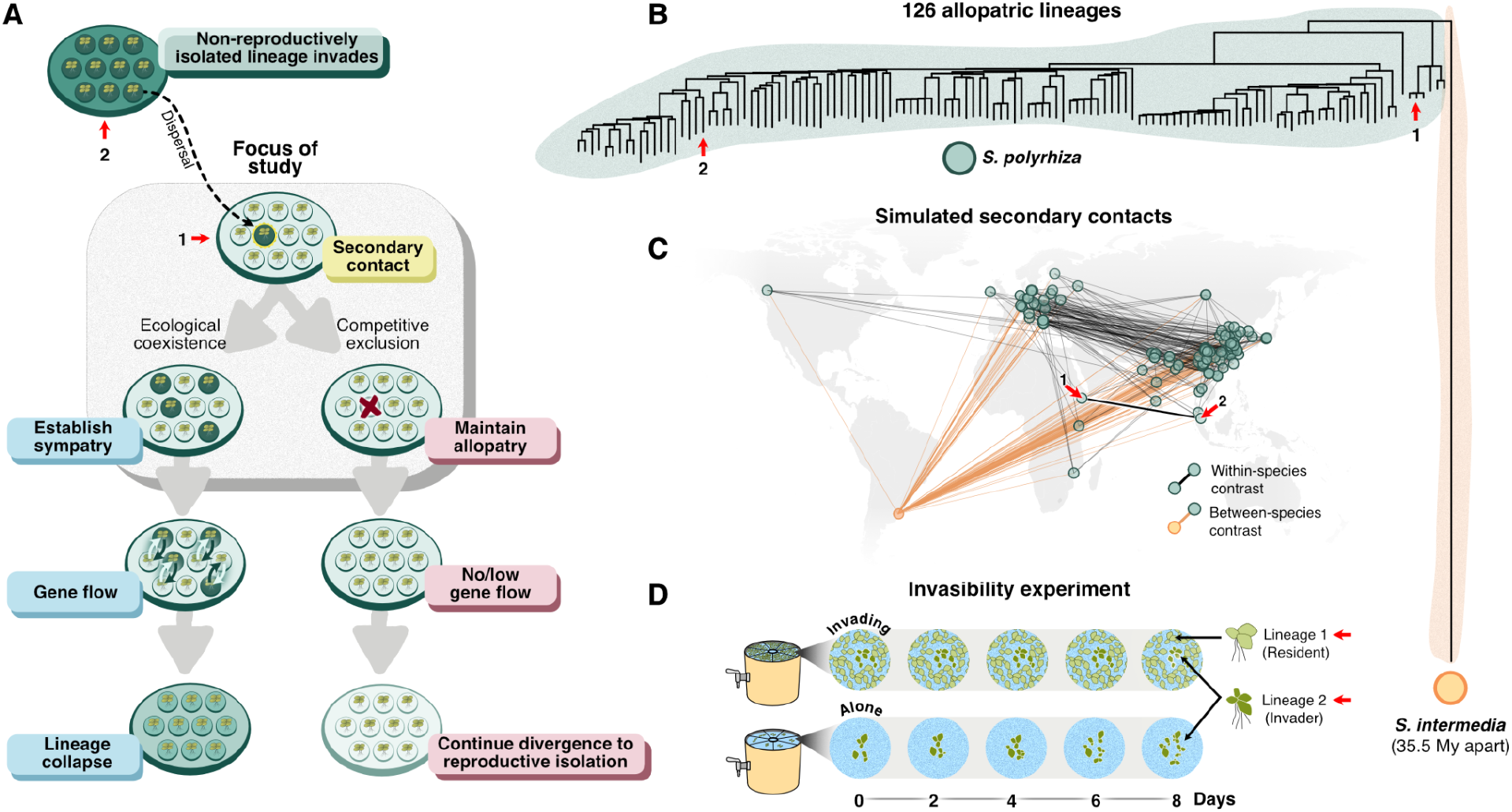
Testing the accumulation of ecological differentiation and coexistence across allopatric and genetically diverging lineages. (**A**) Ecological differentiation can have contrasting consequences for lineage coexistence and speciation following secondary contact. The evolution of greater niche differences among diverging lineages increases the potential for their coexistence in sympatry, permitting the maintenance of interbreeding populations and promoting lineage collapse. The evolution of greater competitive differences increases competitive exclusion in sympatry, permitting further divergence if lineages can persist in allopatry. (**B**) Genetic distance was estimated among 126 lineages of *S. polyrhiza* using genome-wide SNP data. Tips “1” and “2” illustrate two allopatric lineages as an example across panels. The orange circle represents the sister-species *Spirodela intermedia*, which diverged from *S. polyrhiza* ~35.5 mya ^32^. (**C**) Each green point corresponds to a unique *S. polyrhiza* lineage, with black lines denoting pairwise lineages that were competed to test for coexistence in sympatry. To examine ecological differentiation and coexistence beyond the species boundary, we sourced one reference lineage of *S. intermedia* from Uruguay as an outgroup, that we competed against all 126 *S. polyrhiza* lineages (orange lines). (**D**) Competition trials simulating secondary contact were conducted in a common garden within experimental arenas containing recreated pond media. Pairwise coexistence and ecological differentiation was quantified via reciprocal invasion trials, in which the ability of each lineage to grow from rare (*N* = 6 starting individuals of each ‘invader’ lineage) was estimated over 8 days, with and without a competitor ‘resident’ lineage at its carrying capacity. See Supplementary Materials for experimental details.

For mode, modern coexistence theory ^21–23^ states that ecological differentiation takes two opposing forms that together dictate whether diverging lineages have the potential to coexist in sympatry or will be competitively excluded upon secondary contact (Fig. 1A). On one hand, the accumulation of niche differences (e.g., via divergent selection to unique resource niches ^24^) will increase self-limitation more than competition among lineages. As a result, greater niche differences will increase the potential for nascent lineages to coexist in sympatry ^22,25^. By contrast, the accumulation of competitive differences among lineages (e.g., via directional selection to optimize resource-use efficiency ^24^) will increase asymmetry in competitive ability and destabilize coexistence, preventing sympatry upon secondary contact ^22,25^. Thus, if the relative accumulation of competitive differences exceeds that of niche differences, competitive exclusion may delay sympatry until sufficient reproductive isolation has accrued, provided lineages persist in allopatry (Fig. 1A). While the importance of ecological differentiation for speciation has been long recognized (e.g., ^26,27^), whether this differentiation confers a niche or a competitive difference is rarely considered (but see ^17^) despite opposing effects on achieving sympatry during speciation ^11^

Here, we test the mode and tempo by which the potential to coexist upon secondary contact evolves among nascent lineages at evolutionary timescales critical to speciation. We measured coexistence outcomes (henceforth ‘coexistence potential’) across 315 pairings within 126 allopatric, genetically divergent lineages of the common duckweed *Spirodela polyrhiza* – a floating freshwater angiosperm – totalling 1890 competition trials simulating secondary contact in a common environment (Fig. 1). We leveraged the global distribution of *S. polyrhiza* and selected allopatric lineages spanning five continents, with geographic distances between lineages ranging from 50 km to 8,800 km (Fig. 1C) and lineage pairs varying in their degree of genetic divergence (Fig. 1B), which we estimate represents between 9,000 to 68,000 years of geographic separation (see Supplementary Materials for estimation of divergence time). We chose lineages to minimize correlations between genetic distance (a proxy for time ^26^) and spatial distance (fig. S1A and table S1; see Supplementary Materials for details), as the two variables are often correlated, with the latter potentially confounding relationships with the former.

The potential for lineages to coexist in sympatry was quantified using pairwise reciprocal invasion trials ^25^, where stable coexistence occurs when each lineage can grow from rare when invading a competitor lineage at its carrying capacity (i.e., ‘mutual invasibility’) (Fig. 1D; fig. S2). In total, we tracked 117,722 individual plants growing from rare. We then used these invasion growth rates to estimate the sensitivity of each lineage to competition, from which we quantified niche and competitive differences ^25,28^ The advantage of this approach is that it provides a standardized demographic measure of ecological differentiation and its contribution to persistence without needing to identify the often numerous and interacting biological mechanisms responsible for those outcomes ^29^. Notably, although sexual reproduction in *S. polyrhiza* is estimated to be fairly common in nature at the population level ^30,31^, plants reproduce clonally at short experimental timescales. This allows us to quantify invasion growth rates independently for each competing lineage in our experiment, without obfuscation from reproductive interactions like reinforcement and assortative mating ^15^ (see Supplementary Materials for detailed experimental methods).

### Time in allopatry increases the potential to coexist upon secondary contact

We first hypothesized that ecological differentiation accumulates over evolutionary time separating allopatric lineages, for example, as a product of selection to unique environments. We predicted that: (1) both niche differences and competitive differences increase with genetic distance separating isolated lineages; and that (2) this could alter the potential of diverging lineages to coexist upon secondary contact, depending on the relative magnitude of change in niche and competitive differences.

We found that coexistence potential accumulates with the pairwise genetic distance of lineages (β = 1.262, 95% CI: 0.394 to 2.130, *p* = 0.004; Fig. 2A; table S2). Across genetic distances corresponding to divergence times less than 68,000 years (fig S3), we observed coexistence potential to increase by ~15% (from 1.02, 95% CI: 0.87-1.17 to 1.17, 95% CI: 1.03-1.32). Further, we found that higher coexistence potential between diverging lineages was accompanied by a modest but significant increase in niche differences over genetic distance (β = 1.058, 95% CI: 0.397 to 1.718, *p* = 0.001; Fig. 2B; table S2). By contrast, we found that competitive differences did not change significantly over genetic distance (β = 0.835, 95% CI: −2.196 to 3.867, *p* = 0.581; Fig. 2C; table S2). The potential for lineages to coexist therefore emerges before the evolution of reproductive isolation ^30,31^ consistent with a scenario in which sympatry may then allow for hybridization and the collapse of sympatric lineages upon secondary contact (Fig. 1A). Despite observing substantial unexplained variation (adjusted R^2^ = 0.12), we note such variation may be unsurprising as we are quantifying slight ecological differences among closely related genotypes. Across this variation, we were able to detect a modest directional effect of increasing coexistence potential over exceptionally short divergence times (on the order of tens of thousands of years) compared to classic studies of coexistence between species separated by millions of years ^18,20,33^

**Fig. 2.**
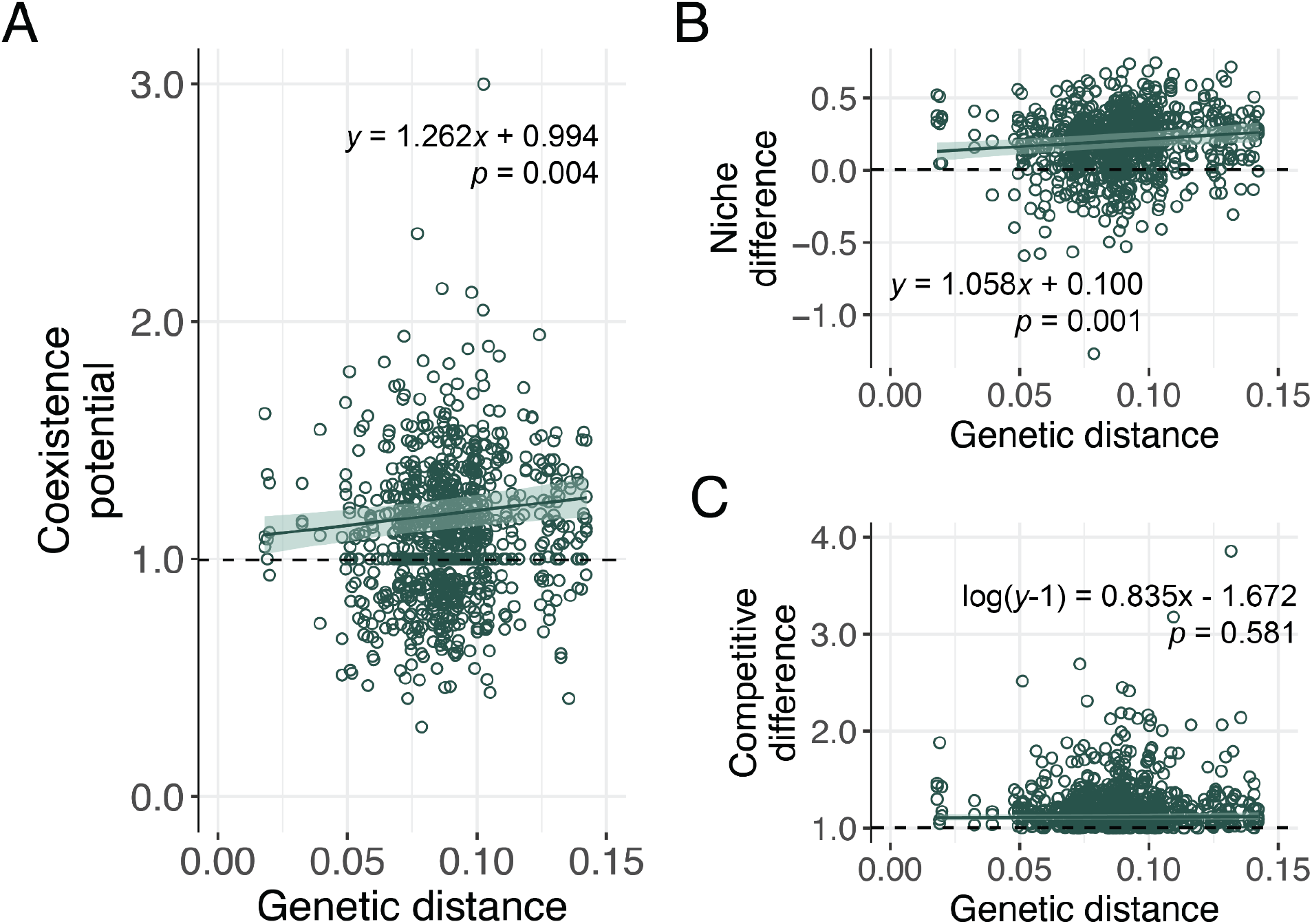
Accumulation of niche differences, competitive differences, and coexistence potential across genetically divergent lineages of *S. polyrhiza*. (**A**) Coexistence potential is the balance of niche and competitive differences (eq. 1.5 in Supplementary Materials), where higher values translate to greater capacity for mutual invasion and pairwise coexistence under experimental conditions. Points above the horizontal, dashed line (>1) represent replicate pairs of *S. polyrhiza* lineages predicted to coexist upon secondary contact, whilst those <1 are predicted to lead to the exclusion of the competitively inferior lineage. **(B)** Higher values above the dotted line (>0) reflect increasing pairwise niche differences. Note that points <0 reflect priority effects (i.e., mutual non-invasibility ^28^). **(C)** Higher values reflect increasing pairwise competitive differences (i.e., greater asymmetry in competitive ability). Note that competitive differences use a shifted response (*y*-1) and are modelled using a Tweedie distribution with a log link (see Supplementary Materials for details). Model predicted slopes (dark green line) and 95% confidence intervals (bands) show that both niche differences and coexistence potential significantly increases over genetic distance. *P* values associated with the slope term are shown. Note that genetic distance represents the proportion of nucleotide differences between lineages (substitutions per site; see Supplementary Materials for details).

Because trait comparisons among pairs of lineages are inherently non-independent ^34,35^, we also conducted a sensitivity analysis using recently developed Bayesian phylogenetic generalized least squares models ^36^ to account for pairwise phylogenetic covariance (see Supplementary Materials for model details). These models recovered similar qualitative patterns, albeit with weaker effect sizes. Specifically, 92.3% of posterior slopes showed a positive relationship between coexistence potential and genetic distance (mean β = 0.021, 95% Highest Posterior Density Interval (HDPI): −0.009 to 0.048; fig. S4 and S5). Niche differences showed a similar trend (89.2% positive slopes; mean β = 0.014, 95% HDPI: −0.008 to 0.035), while evidence of a directional relationship with genetic distance was again weak for competitive differences (64.5% positive slopes; mean β = 0.003, 95% HDPI: −0.013 to 0.019; fig. S4 and S5). We treat these as supporting evidence rather than primary analyses because these phylogenetic models assume a hierarchical and bifurcating phylogeny, whereas genetic relationships among lineages within species are best described by reticulate networks arising from recombination and shared ancestry ^37,38^. Methods that account for pairwise non-independence under such interconnected genealogies do not yet exist.

### Most niche differences evolve prior to the species boundary

The early accumulation of niche differences among nascent lineages prompts two questions: (1) whether niche differences, instead of competitive differences, continue to accumulate beyond microevolutionary timescales within species to macroevolutionary timescales among species ^39^ and (2) whether most niche differences, and therefore coexistence potential, are largely established before speciation is complete (i.e., prior to the species boundary) ^40^. To address this, we conducted an additional 488 competition trials to parameterize niche and competitive differences between each lineage of *S. polyrhiza* to an outgroup lineage of its sister species *Spirodela intermedia*. This allowed us to compare all *S. polyrhiza* lineages against the same standardized competitor (“phytometer”) and quantify pairwise estimates of genetic distance to the *S. intermedia* reference, thereby extending our comparisons across ~35.5 million years of evolutionary divergence^32^ (Fig. 3).

**Fig. 3.**
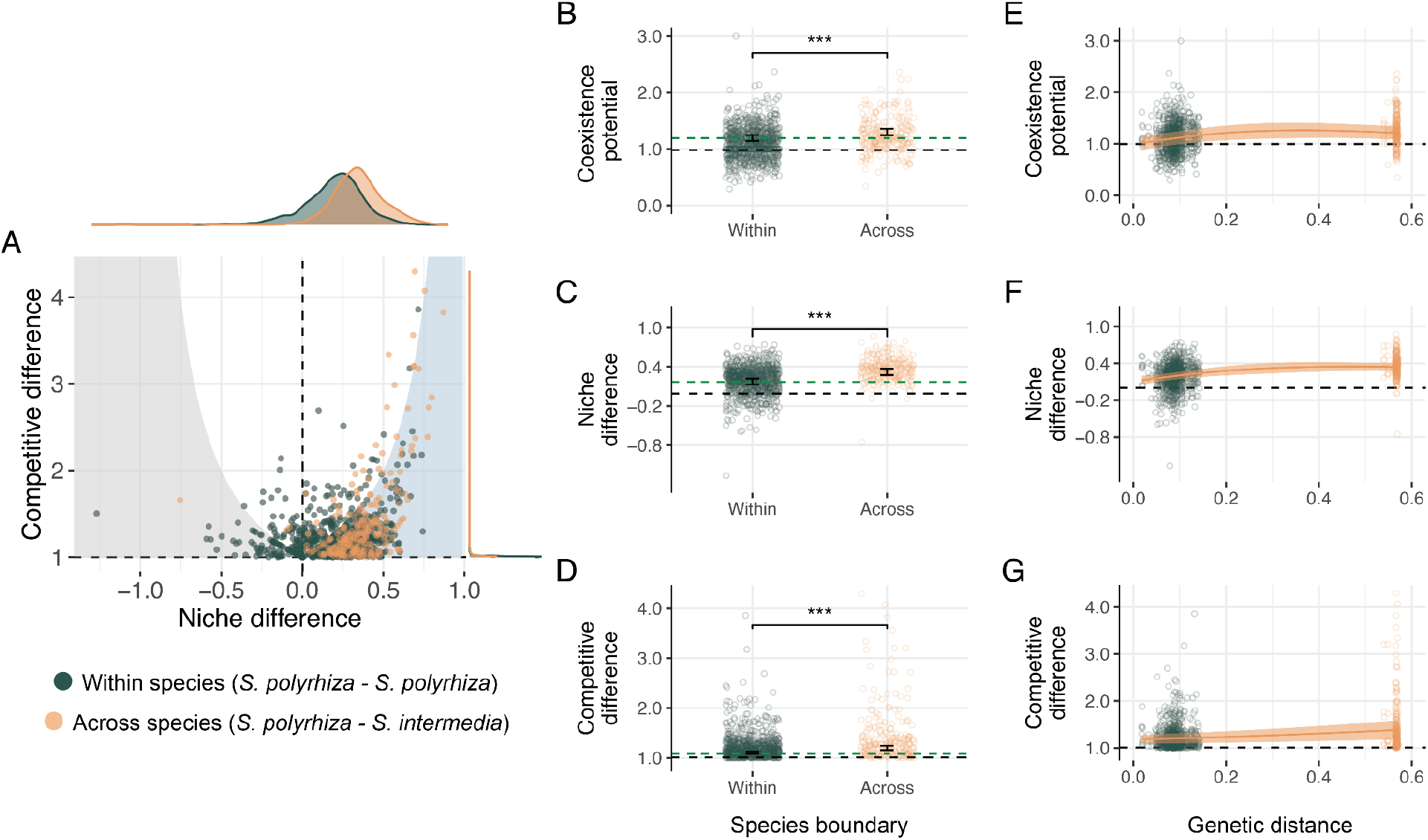
The trajectory of coexistence potential from within to across the species boundary. (**A**) Data points represent pairwise estimates of niche and competitive differences plotted relative to the zone of coexistence (blue shading). Points represent replicate pairs of *S. polyrhiza* lineages competing with other *S. polyrhiza* lineages (dark green) or with their sister species *S. intermedia* (orange). Note that the light gray shading indicates priority effects (niche differences < 0; (*25*)). (**B-D**) Mean differences within and across the species boundary, where *** *p* < 0.001. Error bars represent the 95% confidence interval. Green dashed lines visualize the mean estimate of coexistence potential, niche difference, and competitive difference for within the species boundary. (**E-G**) Trajectory of coexistence potential, niche differences, and competitive differences across genetic distance from within to across the species boundary. Orange trend lines and bands show the predicted mean and 95% CIs from best fit models (i.e., lowest AIC values) when including *S. intermedia* data. Note that genetic distance represents the proportion of nucleotide differences between lineages (substitutions per site; see Supplementary Materials for details).

We found that coexistence potential continued to increase across the species boundary, with greater coexistence across species (i.e., among reproductively isolated lineages) than within species (*β*_*across*_ = 0.106, 95% CI: 0.065 to 0.147, *p* < 0.001; Fig. 3B; table S3). This was also accompanied by larger niche differences across than within species (*β*_*across*_ = 0.149, 95% CI: 0.120 to 0.179, *p* < 0.001; Fig. 3C; table S3). Competitive differences also increased from within to across the species boundary (*β*_a_ _*cross*_ = 0.641, 95% CI: 0.508 to 0.774 on the log-link scale, *p* < 0.001; Fig. 3D; table S3) with *S. intermedia* being competitively dominant over *S. polyrhiza* in 78.7% of all competition trials (χ^2^ = 80.33, *p* < 0.001; fig. S6), although this increase in competitive difference was not enough to shift coexistence outcomes overall. Notably, despite on average greater coexistence potential across than within species, estimates of niche differences, competitive differences, and coexistence showed substantial overlap between and within species (Fig. 3A). We found that 89.8% of between-species estimates for coexistence potential fell within the 95% quantile range of within-species estimates (86.5% and 86.1% overlap for estimates of niche and competitive differences, respectively), suggesting that substantial ecological differentiation that permits coexistence readily exists among competed lineages of *S. polyrhiza*

Furthermore, when between-species contrasts are included, we found that model selection favoured a moderately non-linear and saturating relationship between genetic distance and coexistence potential (AIC = 254.61; Akaike weight (*w*) = 0.885; table S4) over a linear relationship (AIC = 258.69; Akaike *w* = 0.115; table S4), with the fitted smooth term showing modest curvature (EDF = 1.825; Fig. 3E). The slower pace of increase in coexistence potential near the species boundary was also accompanied by a non-linear increase in niche differences (AIC = −513.02; Akaike *w* = 0.758; table S4), which was favoured over a linear increase (AIC = −510.74; Akaike *w* = 0.242; table S4) with genetic distance (EDF = 1.812; Fig. 3F). In contrast, both linear (AIC = −899.79; Akaike *w* = 0.502; table S4) and non-linear models (AIC = −899.78; Akaike *w* = 0.498; table S4) were similarly favoured for changes in competitive difference across species (EDF = 1.000; Fig. 3G). We note that the gap in genetic distance between the two species (Fig. 3E-G) likely reflects both our sampling design, which included only a single lineage of *S. intermedia*, and natural genetic discontinuity between the two reproductively isolated species.

The fitted non-linear relationships should therefore not be interpreted as precise estimates of the underlying evolutionary trajectory in the unsampled region. Nonetheless, our data suggests that ecological differentiation that could permit coexistence in sympatry arises early during lineage divergence and may slow near the speciation boundary.

## Discussion

Darwin predicted that closely related species should compete more intensely and are less likely to coexist in sympatry than distant relatives ^41^. Our results show that this principle extends to much shorter timescales, where the potential for populations to coexist in sympatry increases as they become genetically distinct. Angiosperms are estimated to take ~2.7 million years in allopatry to accrue the genetic incompatibilities required for complete reproductive isolation ^42^ By comparison, we found that in *S. polyrhiza* niche differences that stabilize coexistence accumulated among allopatric lineages that have been separated for no more than 68,000 years. Both the accumulation of niche differences and coexistence potential among genetically diverging lineages (Fig. 2) and their considerable magnitude within versus between species (Fig. 3A) could have key consequences for the outcomes of secondary contact, and thus, patterns of speciation and diversification ^4–7^

First, before the establishment of reproductive isolation, the early evolution of niche differences over competitive differences among allopatric lineages could increase opportunities for interbreeding by permitting their persistence upon secondary contact (Fig. 1A). Early and increasing niche differences among nascent lineages could therefore promote frequent lineage collapse and contribute to time lags in speciation rates ^42^, helping to reconcile the long-debated ‘paradox of stasis’ ^43^ – the common observation that new species tend to arise slowly despite rapid evolution on microevolutionary timescales. Indeed, while *Spirodela* are basal to all other duckweed species this genus has only two extant species (*S. polyrhiza* and *S. intermedia*) over its 35.5 million year history ^32^, consistent with slow divergence rates. While the directional increase in niche differences and coexistence across genetic distance is modest (Fig. 2), we note that this may still translate into a substantial selection differential among populations.

Second, we found that many lineages within species are nearly as ecologically distinct from one another as they are from their sister species (Fig. 3A), and that the accumulation of niche differences and coexistence potential may slow near the species boundary (Fig. 3E and 3F). While this means lineages that establish immediate reproductive isolation –for example in the case of polyploid speciation where without quick ecological differentiation competing lineages may not persist ^11^ –may already be capable of coexisting in sympatry, opportunities for further niche differentiation and diversification may become constrained at broader phylogenetic scales ^5,44^. Furthermore, these results help explain why niche and competitive differences are often non-zero even among very closely related species ^18–20^ and suggests that empiricists should refrain from assuming an absence of differences within species (i.e., by forcing the relationship between ecological differences and phylogenetic distance through the origin ^18^).

Our empirical approach based on niche and competitive differences allows us to avoid exploratory tests of many individual trait differences ^29^ a logistical necessity at the scale of this experiment. Nonetheless, this presents some opportunities for future study, including identifying the trait mechanisms and selective drivers underlying ecological differentiation. To explore one potential driver, we additionally tested whether ecological differences among nascent *S polyrhiza* lineages could be explained by climatic differences a key agent of ecological divergence ^45^–using current climate at the home site of each lineage as a proxy of the selective environment experienced in allopatry. We could not detect an association between climatic dissimilarity and niche differences, competitive differences, or coexistence potential (fig. S7 and table S5; see Supplementary Materials for model details). Genetic distance, however, remained associated with niche differences and coexistence potential after accounting for contemporary climatic dissimilarity (table S5). Together, these results suggest that our climate proxy does not sufficiently capture the historical selective environment, or that other factors such as localized drivers (i.e., resource availability within sites) or drift ^46^ may be at play. While beyond the scope of this study, future studies should test these alternative drivers of niche and competitive differences (e.g., by parameterizing resource-competition models ^24^).

Understanding ecological differentiation among nascent lineages and its consequences for coexistence will be critical for understanding speciation dynamics. By applying coexistence theory across the speciation continuum, our experiment provides a bridge to connect demographic outcomes of ecological divergence at the population level to larger scale patterns of biodiversity ^11^. Our results open several avenues that can be further extended to test the eco-evolutionary mechanisms underlying coexistence prior to speciation, how different phylogenetic patterns emerge in the evolutionary record (e.g., long versus short branches or geographic/latitudinal gradients in diversification rates ^47,48^), and how ecological differentiation interacts with reproductive isolation ^8,17,49^. Together, these tests will advance our understanding of how coexisting species originate ^11^. By uniting ecological and evolutionary perspectives on biodiversity, our results highlight that the processes that generate and maintain biodiversity are tightly linked.

## Acknowledgements

We thank K. Chute, S. Deng, G. Gillies, A. Jackman, L. McBurnie, W. Ou, B. Ozkan, T. Read, C. Spangenberg, M. Tan, and A. Wong for their valuable assistance in setting up the experiment. We are grateful to the staff at UBC South Campus greenhouse for their continued support. We thank M. Sarazova for maintaining and preparing duckweed lines to send to UBC. We thank J. Davies, M. Frederickson, J. Fox, S. Otto, R. Sargeant, D. Schluter, M. Tseng, and J. Williams for comments on earlier versions of the manuscript.

## Author Contributions

Conceptualization: ALA, JS, MMT, RMG, SPH, TU.

Methodology: JS, TU, PD.

Investigation: JS, TU.

Visualization: JS, TU.

Funding acquisition: ALA, RMG.

Project administration: JS, TU.

Supervision: ALA, RMG.

Writing - original draft: ALA, JS, RMG, TU.

Writing - review & editing: ALA, JS, MMT, PD, RMG, SPH, SX, TU.

## Data and materials availability

All data and code is available on Zenodo. DOI: 10.5281/zenodo.23066147

## Supplementary Materials

Materials and Methods

Supplementary Text

Figs. S1 to S12

Tables S1 to S8

References (*1-77*)

## References and Notes

1. Dynesius, M. & Jansson, R. Persistence of within-species lineages: a neglected control of speciation rates: perspective. Evolution 68, 923–934 (2014).

2. Harvey, M. G., Singhal, S. & Rabosky, D. L. Beyond reproductive isolation: demographic controls on the speciation process. Annu. Rev. Ecol. Evol. Syst. 50, 75–95 (2019).

3. Weber, M. G. & Strauss, S. Y. Coexistence in close relatives: beyond competition and reproductive isolation in sister taxa. Annu. Rev Ecol. Evol. Syst. 47, 359–381 (2016).

4. Rabosky, D. L. Diversity-dependence, ecological speciation, and the role of competition in macroevolution. Annu. Rev. Ecol. Evol. Syst. 44, 481–502 (2013).

5. Price, T. D. et al. Niche filling slows the diversification of Himalayan songbirds. Nature 509 222–225 (2014).

6. Mayrose, I. et al. Recently formed polyploid plants diversify at lower rates. Science 333 1257–1257 (2011).

7. Schluter, D. Speciation, ecological opportunity, and latitude Am. Nat 187, 1–18 (2016).

8. Lerch, B. A., Bürger, R. & Servedio, M. R. Reconciling Santa Rosalia: both reproductive isolation and coexistence constrain diversification. Am. Nat. 204, E99–E114 (2024).

9. Januario, M., Pinsky, M. L. & Rabosky, D. L. The metapopulation bridge to macroevolutionary speciation rates: a conceptual framework and empirical test. Ecol. Lett 28, e70021 (2025).

10. Schluter, D. Variable success in linking micro- and macroevolution. Evol. J. Linn. Soc. 3 kzae016 (2024).

11. Germain, R. M. et al. On the origin of coexisting species. Trends Ecol. Evol. 36, 284–293 (2021).

12. Ricklefs, R. E. & Bermingham, E. The causes of evolutionary radiations in archipelagoes: Passerine birds in the Lesser Antilles. Am. Nat 169 285–297 (2007).

13. Rosenblum, E. B. et al. Goldilocks meets Santa Rosalia: an ephemeral speciation model explains patterns of diversification across time scales. Evol. Biol. 39, 255–261 (2012).

14. Schluter, D. The Ecology of Adaptive Radiation (OUP Oxford, 2000).

15. Rundle, H. D. & Nosil, P. Ecological speciation. Ecol. Lett. 8, 336–352 (2005).

16. Baack, E., Melo, M. C., Rieseberg, L. H. & Ortiz-Barrientos, D. The origins of reproductive isolation in plants. New Phytol 207, 968–984 (2015).

17. Irwin, D. & Schluter, D. Hybridization and the coexistence of species. Am. Nat 200 E93–E109 (2022).

18. Godoy, O., Kraft, N. J. B. & Levine, J. M. Phylogenetic relatedness and the determinants of competitive outcomes. Ecol. Lett. 17, 836–844 (2014).

19. Narwani, A., Alexandrou, M. A., Oakley, T. H., Carroll, I. T. & Cardinale, B. J. Experimental evidence that evolutionary relatedness does not affect the ecological mechanisms of coexistence in freshwater green algae. Ecol. Lett 16, 1373–1381 (2013).

20. Germain, R. M., Weir, J. T. & Gilbert, B. Species coexistence: macroevolutionary relationships and the contingency of historical interactions. Proc. R. Soc. B Biol. Sci 283 20160047 (2016).

21. Chesson, P. MacArthur’s consumer-resource model. Theor. Popul. Biol. 37, 26–38 (1990).

22. Chesson, P. Mechanisms of maintenance of species diversity. Annu. Rev. Ecol. Evol. Syst. 31 343–366 (2000).

23. Chesson, P. & Kuang, J. J. The interaction between predation and competition. Nature 456 235–238 (2008).

24. Sakarchi, J. MacArthur’s consumer-resource model: a Rosetta Stone for competitive interactions. Am. Nat. 205, 306–326 (2025).

25. Carroll, I. T., Cardinale, B. J. & Nisbet, R. M. Niche and fitness differences relate the maintenance of diversity to ecosystem function. Ecology 92, 1157–1165 (2011).

26. Funk, D. J., Nosil, P. & Etges, W. J. Ecological divergence exhibits consistently positive associations with reproductive isolation across disparate taxa. Proc. Natl. Acad. Sci. 103 3209–3213 (2006).

27. Anacker, B. L. & Strauss, S. Y. The geography and ecology of plant speciation: range overlap and niche divergence in sister species. Proc R. Soc. B Biol. Sci. 281, 20132980 (2014).

28. Grainger, T. N., Levine, J. M. & Gilbert, B. The invasion criterion: a common currency for ecological research. Trends Ecol. Evol. 34, 925–935 (2019).

29. Germain, R. M., Williams, J. L., Schluter, D. & Angert, A. L. Moving character displacement beyond characters using contemporary coexistence theory. Trends Ecol. Evol 33, 74–84 (2018).

30. Xu, S. et al. Low genetic variation is associated with low mutation rate in the giant duckweed. Nat. Commun. 10, 1243 (2019).

31. Wang, Y. et al. Population genomics and epigenomics of Spirodela polyrhiza provide insights into the evolution of facultative asexuality. Commun. Biol 7, 1–12 (2024).

32. Tippery, N. P. & Les, D. H. Tiny plants with enormous potential: Phylogeny and evolution of duckweeds. The duckweed genomes (2020).

33. Cadotte, M. W., Davies, T. J. & Peres-Neto, P. R. Why phylogenies do not always predict ecological differences. Ecol. Monogr. 87, 535–551 (2017).

34. Fitzpatrick, B. M. Molecular correlates of reproductive isolation Evolution 56, 191–198 (2002).

35. Castillo, D. M. Factors contributing to the accumulation of reproductive isolation: A mixed model approach. Ecol. Evol 7, 5808–5820 (2017).

36. Anderson, S. A. S., Kaushik, S. & Matute, D. R. The comparative analysis of lineage-pair traits. BioRxiv Prepr. Serv. Biol. 2024.11.28.625927 (2024) doi:10.1101/2024.11.28.625927.

37. Maddison, W. P. Gene trees in species trees. Syst. Biol. 46, 523–536 (1997).

38. Maddison, W. P. & Knowles, L. L. Inferring phylogeny despite incomplete lineage sorting. Syst. Biol. 55, 21–30 (2006).

39. Rolland, J. et al. Conceptual and empirical bridges between micro- and macroevolution. Nat. Ecol. Evol 7, 1181–1193 (2023).

40. Rundell, R. J. & Price, T. D. Adaptive radiation, nonadaptive radiation, ecological speciation and nonecological speciation. Trends Ecol. Evol. 24 394–399 (2009).

41. Darwin, C. On the Origin of Species by Means of Natural Selection, or the Preservation of Favoured Races in the Struggle for Life. (John Murray, United Kingdom of Great Britain and Ireland, 1859).

42. Hedges, S. B., Marin, J., Suleski, M., Paymer, M. & Kumar, S. Tree of life reveals clock-like speciation and diversification. Mol. Biol. Evol. 32 835–845 (2015).

43. Haller, B. C. & Hendry, A. P. Solving the paradox of stasis: squashed stabilizing selection and the limits of detection. Evolution 68, 483–500 (2014).

44. Ricklefs, R. E. Evolutionary diversification, coevolution between populations and their antagonists, and the filling of niche space. Proc Natl. Acad. Sci. 107, 1265–1272 (2010).

45. Wiens, J. J. The niche, biogeography and species interactions. Philos. Trans. R. Soc. B Biol Sci 366, 2336–2350 (2011).

46. Anderson, S. A. S. & Weir, J. T. The role of divergent ecological adaptation during allopatric speciation in vertebrates. Science 378, 1214–1218 (2022).

47. Schluter, D. & Pennell, M. W. Speciation gradients and the distribution of biodiversity. Nature 546, 48–55 (2017).

48. Weir, J. T. & Lawson, A. Evolutionary rates across gradients. Methods Ecol. Evol 6 1278–1286 (2015).

49. Felsenstein, J. Skepticism towards Santa Rosalia, or why are there so few kinds of animals? Evolution 35, 124–138 (1981).

50. Jacobs, D. L. An ecological life-history of Spirodela polyrhiza (greater duckweed) with emphasis on the turion phase. Ecol. Monogr. 17, 437–469 (1947).

51. Les, D. H., Crawford, D. J., Landolt, E., Gabel, J. D. & Kimball, R. T. Phylogeny and systematics of Lemnaceae, the duckweed family. Syst. Bot 27, 221–240 (2002).

52. Crawford, D., Landolt, E., Les, D. & Kimball, R. Speciation in duckweeds (Lemnaceae): phylogenetic and ecological inferences. Aliso 22 231–242 (2006).

53. Landolt, E. Biosystematic Investigations in the Family of Duckweeds, Lemnaceae: The Family of Lemnaceae, a Monographic Study. Volume 2: Morphology, Karyology, Ecology, Geographic Distribution, Systematic Position, Nomenclature Descriptions. (Geobotanisches Institut der ETH, Zurich, Switzerland, 1986).

54. Bog, M., Appenroth, K. J. & Sree, K. S. Key to the determination of taxa of Lemnaceae: an update. Nord. J. Bot. 38, (2020).

55. Landolt, E. Lemnaceae duckweed family. J. Ariz.-Nev. Acad. Sci 26, 10–14 (1992).

56. Ho, E. K. H., Bartkowska, M., Wright, S. I. & Agrawal, A. F. Population genomics of the facultatively asexual duckweed Spirodela polyrhiza New Phytol. 224, 1361–1371 (2019).

57. Appenroth, K.-J., Teller, S. & Horn, M. Photophysiology of turion formation and germination in Spirodela polyrhiza Biol. Plant 38 95–106 (1996).

58. Docauer, M. A nutrient basis for the distribution of the Lemnaceae. (University of Michigan, Ann Arbor, Michigan).

59. Schneider, C. A., Rasband, W. S. & Eliceiri, K. W. NIH Image to ImageJ: 25 years of image analysis. Nat. Methods 9, 671–675 (2012).

60. Hart, S. P., Freckleton, R. P. & Levine, J. M. How to quantify competitive ability. J. Ecol. 106, 1902–1909 (2018).

61. Gillies, G. J., Angert, A. L. & Usui, T. Temperature dependence and genetic variation in resource acquisition strategies in a model freshwater plant. Funct. Ecol 38, 1600–1610 (2024).

62. Armitage, D. W. & Jones, S. E. Negative frequency-dependent growth underlies the stable coexistence of two cosmopolitan aquatic plants. Ecology 100, e02657 (2019).

63. Spaak, J. W. & De Laender, F. Intuitive and broadly applicable definitions of niche and fitness differences. Ecol. Lett. 23, 1117–1128 (2020).

64. Spaak, J. W., Ke, P.-J., Letten, A. D. & De Laender, F. Different measures of niche and fitness differences tell different tales. Oikos 2023 e09573 (2023).

65. MacArthur, R. Species packing and competitive equilibrium for many species. Theor. Popul. Biol. 1, 1–11 (1970).

66. Grainger, T. N., Letten, A. D., Gilbert, B. & Fukami, T. Applying modern coexistence theory to priority effects. Proc. Natl. Acad. Sci. U. S. A 116, 6205–6210 (2019).

67. Wang, W. et al. The Spirodela polyrhiza genome reveals insights into its neotenous reduction fast growth and aquatic lifestyle. Nat. Commun 5 3311 (2014).

68. Felsenstein, J. Phylogenies from molecular sequences : inference and reliability. Annu. Rev. Genet 22, 521–565 (1988).

69. Paradis, E. & Schliep, K. ape 5.0: an environment for modern phylogenetics and evolutionary analyses in R. Bioinformatics 35, 526–528 (2019).

70. Höhna, S. et al. RevBayes: Bayesian phylogenetic inference using graphical models and an interactive model-specification language. Syst. Biol. 65, 726–736 (2016).

71. RDC Team. R: A language and environment for statistical computing. (2020).

72. Wood, S. N. Fast stable restricted maximum likelihood and marginal likelihood estimation of semiparametric generalized linear models. J. R. Stat. Soc. Ser. B Stat. Methodol. 73, 3–36 (2011).

73. Krishnamoorthy, K. & Lee, M. Improved tests for the equality of normal coefficients of variation. Comput. Stat. 29, 215–232 (2014).

74. Marwick, B. & Krishnamoorthy, K. cvequality: tests for the equality of coefficients of variation from multiple groups. R Softw. Package Version 01 3, (2019).

75. Johnson, J. B. & Omland, K. S. Model selection in ecology and evolution. Trends Ecol. Evol. 19, 101–108 (2004).

76. Armitage, D. W. Global maps of lake surface water temperatures reveal pitfalls of air-for-water substitutions in ecological prediction. Ecography 2023, e06595 (2023).

77. Armitage, D. W. Coexistence barriers confine the poleward range of a globally distributed plant. Ecol. Lett. 23, 1838–1848 (2020).

