## Supplementary Materials for "Niche differentiation confers coexistence prior to the species boundary in an aquatic plant"

**The PDF file includes:**

Materials and Methods
Figs. S1 to S12
Tables S1 to S8
References (*1-77*)

### MATERIALS & METHODS

#### a. *SPIRODELA* COLLECTION AND CULTURING

Duckweeds are free-floating, freshwater angiosperms. We used duckweeds in the *Spirodela* genus (family Araceae)[50,51], which comprises two species (*S. polyrhiza* and *S. intermedia*) that separated around 35.5 mya [32]. *S. polyrhiza* has a near-global distribution while its sister species *S. intermedia* is restricted to warm, temperate climates in central and South America [52]. An individual plant consists of a small, circular frond with many rootlets attached on the underside, with *S. intermedia* having fronds that are smoother, more oblong, and lighter green than *S. polyrhiza* [53,54]. Plants compete for nutrients, space, and light [50,55] and under ideal conditions have a rapid generation time of 3-7 days [53]. Flowering at the individual-level is rare and reproduction occurs exclusively via the asexual budding of clonal, daughter fronds during experiments [50,55]. However, sexual reproduction is estimated to be relatively frequent at the population level, shaping patterns of genetic differentiation among natural populations [30,56].

We used 126 genetically unique and allopatric lineages of *S. polyrhiza* to estimate the potential for coexistence within species. Additionally, we sourced one lineage of *S. intermedia* as an outgroup to estimate coexistence between species. As our main focus was on coexistence within species and across a range of intraspecific genetic distances, we only used a single lineage of *S. intermedia* as a point of reference representing much larger genetic distances. All plants were obtained from a long-term stock collection at the Institute of Organismic and Molecular Evolution, University of Mainz, Germany. We chose *S. polyrhiza* lineages so that experimental lineages originated from a broad geographical range and represented a wide range of pairwise genetic distances (see Fig. 1 in main text). We also chose experimental lineages such that across all lineages any existing correlation between pairwise genetic distance and spatial distance was minimised (fig. S1A). This ensured that pairs of lineages that were more genetically distant were not necessarily lineages that just came from further apart in space. This allowed us to separate the effects of geography and evolutionary time in downstream analyses, and test whether coexistence and its mechanisms change over genetic distance *per se* (see fig. S8 and table S1 for analyses of spatial distance on coexistence).

We propagated and maintained all plants under axenic conditions in a laboratory at the University of British Columbia, Canada, growing each lineage separately in 250 mL Erlenmeyer flasks with 100 mL of artificial pond media (table S6) [57], under constant temperatures (20 °C), and full-spectrum LED lighting (SunBlaster 6400K). Growth medium was refreshed around every 1.5 months to maintain lab stocks of each lineage.

### b. EXPERIMENTAL SETUP AND POPULATION EQUILIBRATION

We conducted pairwise competition trials simulating secondary contact in buckets (23 cm x 28 cm; D x H) placed atop greenhouse benches. Each bucket was filled with 6 L of artificial pond media [58] (table S6) of which 10 % was replaced every two days during the experiment. This was done to ensure chemostat-like conditions that maintained a constant resource supply, a core assumption of competition models that are parameterized to estimate ecological coexistence [24].

To conduct an experimentally tractable number of pairwise competition trials, we first grouped all 126 *S. polyrhiza* lineages into 21 experimental groups ($N = 6$ unique lineages per group) within which we conducted reciprocal competition experiments for all possible pairwise combinations (i.e., $N = 15$ unique pairwise combinations per group). We chose these experimental groups so that correlations between pairwise genetic distance and spatial distance were still minimised within each group ($r = 0.003$ to 0.347). We additionally competed all 126 *S. polyrhiza* lineages with the single *S. intermedia* lineage. Together, we therefore conducted competition trials for 441 unique pairwise combinations of lineages (315 pairwise combinations of *S. polyrhiza* lineages plus 126 pairwise combinations of *S. polyrhiza* – *S. intermedia* lineages).

Before conducting competition trials, we must first grow one competitor lineage ('resident') to its carrying capacity. While the exact value of carrying capacities is not necessary for our methodology (see **d. ESTIMATING NICHE AND COMPETITIVE DIFFERENCES**), it is critical that residents reach a stable equilibrium density such that competitive interactions reflect the equilibrial resource conditions assumed under competitive models used to estimate coexistence. To grow the resident lineage to its equilibrium population density, we initially seeded 200 individuals of each lineage inside each bucket. We tracked each resident's population size by photographing a subsample of each population (1/7th of the total bucket area) and quantifying the percent plant cover every 2-4 days using *ImageJ* (v1.53) [59]. From this data, we tracked resident lineages to a state where their birth rate was approximately equal to their death rate (~3 months; fig. S2) before introducing competing lineages. We note substantial variation in surface area at equilibrium among lineages (fig. S2) suggesting that space is unlikely to be the sole limiting resource of the system. Variation in resident carrying capacity contributes in part to variation in invasion growth rates, because greater resident density lowers resource availability for invading genotypes [24] and because carrying capacity is also a component of the resident's competitive ability [60].

### c. COMPETITION TRIALS

After resident lineages had reached their carrying capacity, we introduced the second competitor lineage ('invader') by adding $N = 6$ individuals (mean = 6.0; SD = 0.7) of each invader lineage within the established residents (fig. S9). For *S. polyrhiza* residents, we simultaneously introduced five *S. polyrhiza* invaders (i.e., the five other lineages that are in the same experimental group as the resident *S. polyrhiza* lineage) plus one *S. intermedia* invader within each bucket (for a total of 6 invaders per bucket x 126 buckets, with one bucket for every *S.*

*polyrhiza* lineage serving as the resident). For *S. intermedia* residents, we simultaneously introduced six *S. polyrhiza* invaders (for a total of 6 invaders per bucket x 21 buckets, with each bucket competing against one of the 21 *S. polyrhiza* experimental groups). We do not expect any interference between invading lineages within each bucket, as each invader: (i) was placed equidistant from each other; and (ii) was seeded at very low densities relative to the density of resident lineages at equilibrium (see fig. S9).

We tracked the growth of each invader lineage (i.e., the 'invasion' growth rate) over 8 days. While this is a relatively short period of time, this is a necessary requirement of invasion trials to satisfy the assumption that the invader remains rare and does not influence their own growth or that of the resident population [28]). Since we were mostly competing lineages within the same duckweed species, we visually tracked the invasion growth rate of each invader by marking each budding frond with a small white dot using acrylic paint every two days using a 200 µL pipette tip. We also tracked mortality by marking invader fronds that were yellowing over the eight days. We note that due to the large number of resident individuals we could not mark resident fronds. In total, we repeated each competition trial across three temporal blocks, with one replicate per block (3 replicates x 6 invaders x 147 buckets = 2646 competition trials).

During these eight days, we also estimated the low-density growth rate of every lineage without competitors (i.e., its 'monoculture' growth rate [28]) using an additional 42 buckets in the greenhouse. We did this by introducing $N = 6$ individuals of each lineage (mean = 6.0; SD = 0.7) into a bucket without any resident lineage, adding 6-7 unique lineages simultaneously within each bucket (for a total of $N = 36$-42 invading individuals per bucket). As a sham-control, we also marked these fronds with acrylic paint every two days to track growth rates. Because both invasion and low-density treatments are marked, we are able to reduce potential systematic bias due to marking when calculating sensitivity in our downstream analysis [1.1]. For each lineage, we repeated these monoculture growth trials across three temporal blocks, with two replicates per block (6 replicates x 127 lineages = 762 monoculture growth trials).

All buckets were rotated 90º every two days during the experiment to reduce light heterogeneity within buckets. Each bucket was also covered with a clear, plastic lid to prevent cross-contamination of nutrient media between buckets. All greenhouse experiments were conducted between June and October 2022. In total, we tracked 117,722 invading plant individuals across all buckets during the experiment. We note that for the competition and monoculture growth trials, we sourced introduced lineages from lab-maintained stocks. To ensure that lab-sourced plants did not have a significant competitive advantage over residents, all invading lineages were first grown in the greenhouse within clear test tubes filled with water for two days prior to the competition trial. This acclimatization period was used to reduce nutrient stores that could have otherwise conferred a competitive advantage for lab-sourced plants [61]. Further, we do not expect systematic differences in invader and resident age, as invaders were sourced from large and mixed-age stock populations. It remains possible that estimated low-density and invasion growth rates may be inflated if, for example, death rates lag beyond what we could observe during the 8-day invasion period, or if there are carry-over effects that our acclimatization period was unable to fully eliminate, although we note that this would not bias slope estimates across genetic distance. Reliably accounting for such an effect would require substantially more replication of lineage-specific baseline measurements (e.g., self-invasions), which our experiment was not designed to provide. Finally, we note that *S. polyrhiza* can form dormant, overwintering turions [32], although the effect of this storage effect on coexistence is found to be relatively weak in *S. polyrhiza* duckweeds competing with another duckweed species, *Lemna minor* [62]. To prevent turions from resurfacing during the experiment and contributing to population dynamics across different temporal blocks and across different invading lineages, we had to remove turions from all invader fronds as soon as they were separated from the mother frond. To prevent bias in low-density growth rates, we also removed and excluded turion counts from lineages introduced without competitors. Nonetheless, we note that the number of turions produced was negligible in our experiment: 93% of censused populations had ≤1 turion across all competition growth trials (92% across all low-density growth trials). Moreover, we did not observe turion number to be biased across the genetic distance of competing lineages (fig. S10).

### d. ESTIMATING NICHE AND COMPETITIVE DIFFERENCES

There are many methods to estimate coexistence and its component mechanisms, though perhaps one of the most robust is the methodology outlined by Carroll et al. [25,63] as it: (i) is not restricted in how competition operates (i.e., is agnostic to whether there is resource competition for biotic resources that follow logistic growth, abiotic resources that follow steady, chemostat-like supply rates, or interference competition); and (ii) is not restricted to any particular model of competitive population dynamics. Moreover, we chose this method a priori as it is particularly suitable for fast growing organisms like duckweed where invasion growth rates can be directly measured [56,57]. While model choice can quantitatively and qualitatively affect estimates of niche and competitive differences, the Carroll method bridges conceptually different approaches used to estimate niche and competitive differences while also producing estimates that are broadly correlated to other commonly used models [64].

The Carroll method is centered on MacArthur's consumer-resource model in continuous time [65], and relies on first estimating the sensitivity to competition for each lineage, from which we can then derive niche differences, competitive differences (also known as 'competitive asymmetry' or ecological 'fitness differences' [22]), and the potential for stable, local coexistence. Sensitivity to competition for lineage $i$ ($S_{ij}$) is estimated as the amount by which its growth rate is reduced when growth is measured in the presence of a competitor lineage $j$ ($g_{ij}$) at its population equilibrium versus when growth is measured alone without the competitor lineage ($g_{i0}$):

$$S_{ij} = \frac{g_{i0} - g_{ij}}{g_{i0}} \qquad [1.1]$$

Where $g_{i0}$ and $g_{ij}$ were estimated with an exponential growth model using individual frond counts obtained every 2 days for 8 days (see **c. COMPETITION TRIALS**). An increase in $S_{ij}$ therefore translates to greater sensitivity of lineage *i* to lineage *j*. Therefore, when $S_{ij} < 1$ then lineage *i* can invade *j*, whereas if $S_{ij} > 1$ then *i* cannot invade *j*. With two competing lineages, we can label the first lineage as the one that is most sensitive to competition ($S_{ij} > S_{ji}$). Because coexistence requires that both lineages can invade when rare (all $g_{ij} > 0$), we must then have $1 > S_{ij}$ for stable coexistence.

The Carroll method [25] uses these sensitivities to define niche ($ND_{(i,j)}$) and competitive differences ($CD_{(i,j)}$) between each pair of competitors. Stabilising niche differences are estimated as 1 minus the niche overlap ($\rho_{(i,j)}$), where $\rho_{(i,j)}$ is calculated as the geometric mean of lineage *i* and *j*'s sensitivities:

$$\rho_{(i,j)} = \sqrt{S_{ij} S_{ji}} \qquad [1.2]$$

$$ND_{(i,j)} = 1 - \rho_{(i,j)} \qquad [1.3]$$

Therefore, if on average a pair of lineages are less sensitive to competition there would be reduced niche overlap, and therefore increased niche differences, between these lineages. On the other hand, competitive differences ($CD_{(i,j)}$) are estimated as the geometric standard deviation of lineage *i* and *j*'s sensitivities:

$$CD_{(i,j)} = \sqrt{S_{ij}/S_{ji}} \qquad [1.4]$$

Therefore, greater variability in the sensitivities (i.e., one lineage is more sensitive to competition than the other) would mean increased competitive differences between these lineages.

Whether coexistence or competitive exclusion occurs for a given pair of competing lineages depends on the relative balance of niche and competitive differences [22]. As stated above, stable coexistence is achieved when the most sensitive lineage remains able to invade the second lineage at its population equilibrium ($1 > S_{ij}$). Narwani et al. [19] stated that this coexistence criterion is equivalent to the following condition:

$$\frac{1}{CD_{max}(1 - ND_{(i,j)})} > 1 \qquad [1.5]$$

where $CD_{max}$ represents the larger of the two reciprocal competitive differences ($CD_{(i,j)}$ or $CD_{(j,i)}$). That is, niche differences must be sufficiently large and competitive differences low enough for both lineages to coexist. We use equation [1.5] as our metric of coexistence potential in sympatry, such that values greater than 1 correspond to increasing potential for coexistence. We note coexistence potential [1.5] can also be written in terms of sensitivities, where coexistence potential represents the reciprocal of the most sensitive invader ($|S_{ij}^{-1}|$ or $|S_{ji}^{-1}|$). Thus, when the more sensitive competitor's invasion growth rate equals zero, using equation [1.1], coexistence potential takes the value of exactly one. We also note that priority effects are achieved when both lineages are unable to invade an already established competitor. This occurs when both $S_{ij}$ and $S_{ji}$ are > 1, or if expressed in terms of niche and competitive differences, when $ND < 0$ and $CD < 1 - ND$ [66]. Finally, we note that negative sensitivity values correspond to cases of facilitation [25], which can lead to complex values when estimating competitive differences [1.4]. However, facilitation only represented 0.5% of all competition trial outcomes, and so such cases were dropped from all downstream analyses.

### e. ESTIMATING GENETIC DISTANCE

Whole genome sequences of all 126 *S. polyrhiza* lineages were obtained and sequenced as part of a prior study on the population genomics of *S. polyrhiza* [30,31]. We additionally obtained the whole genome sequence of one *S. intermedia* lineage for our experiment. Briefly, sequences were obtained using Illumina short-read sequencing with 150 bp paired-end reads at ~29X coverage. All sequence reads were then trimmed, aligned to the *S. polyrhiza* reference genome [67], checked for mapping quality, then called and filtered for genetic variants (SNPs and small indels). From this genome-wide SNP data, we created a pairwise genetic distance matrix among experimental lineages using PHYLIP v3.69 [68] and constructed a neighbor-joining tree using the *ape* package (version 5.8; [69]) in R (see Fig. 1 in in main text).

We additionally obtained a range of divergence time estimates among *S. polyrhiza* lineages in our experiment to estimate the timescale at which niche and competitive differences evolved (fig. S3). We estimated divergence times using RevBayes (version 1.2.1; [70]). Specifically, we ran a Bayesian coalescent analysis to estimate split times at each node in the genealogy of *S. polyrhiza* using *S. intermedia* as an outgroup. We implemented the *dnCoalescent* function, which models a constant coalescent process to infer the genealogy [70]. A molecular clock was also implemented using a secondary calibration for the root age of *S. polyrhiza* and *S. intermedia* set to 35.5 mya [32]. We set a range around this root age using a normal distribution where SD = 6 my, minimum = root age – 12.5 my, and maximum = root age + 12.5 my. To reduce computational time, we split our 126 *S. polyrhiza* lineages into 4 subgroups and estimated divergence time for each subgroup. We ran each MCMC model for 20,000 iterations, thinning every 10th iteration. We used the default, uniform prior for the population size with constant lower and upper bounds set to 0 and $1 \times 10^8$, respectively.

### f. MODEL FITTING

All model-fitting was conducted in R (version 4.3.0; [71]). We fit all models below as a generalized additive mixed-effects model (GAMM) using the *mgcv* package (version 1.9.1; [72]), with smoothing parameters estimated by restricted maximum likelihood. All models included temporal block and experimental group as random effects, with the latter accounting for pairwise estimates being obtained from the same set of buckets within an experimental group. Since competitive differences were positively skewed with a lower bound of 1, we applied a shift of -1 to place the lower bound at zero and used a Tweedie distribution modelled on a log-link scale for downstream models.

We first tested how niche differences, competitive differences, and the potential for coexistence varied across genetically diverging *S. polyrhiza* lineages. We fit a separate model for each response variable with a linear effect of pairwise genetic distance as the fixed predictor (Fig. 2 in main text; table S2). We then tested if coexistence and its mechanisms continued to change from within species to across the species boundary by including competition data from *S. polyrhiza–S. intermedia* pairs. We tested if mean estimates of niche differences, competitive differences, and coexistence differed within (i.e., *S. polyrhiza–S. polyrhiza* pairs) compared to across the species boundary (i.e., *S. polyrhiza–S. intermedia* pairs), by fitting species level as a categorical predictor variable (table S3; Fig. 3B-D in main text). We additionally tested if variability in estimates of niche differences, competitive differences, and coexistence differed within compared to across the species boundary (fig. S11; table S7). To allow for statistical comparisons of variability, we first estimated the coefficient of variation (CV) separately across *S. polyrhiza–S. polyrhiza* pairs or *S. polyrhiza–S. intermedia* pairs for each of the 21 experimental groups. Then, we tested for differences in CV within versus between the species boundary by conducting a Modified signed-likelihood ratio test (MLRT) for the equality of CVs, which accounts for uneven sample sizes and is more robust to type I error [73]. We ran tests on equality of CVs using the package *cvequality* (v0.1.3; [74]).

Lastly, to understand how niche and competitive differences change from within to across the species boundary, we included data from competition trials between *S. polyrhiza* and *S. intermedia*. We fit both linear and nonlinear effects of pairwise genetic distance, with the latter being modelled using a spline with a restricted basis dimension ($k = 3$) to avoid over-fitting (note that for competitive differences linear and nonlinear effects were evaluated on the log-link scale). We then used the Akaike Information Criterion (AIC) to assess whether the linear or nonlinear model better described the trajectory of niche differences, competitive differences, and coexistence across the species boundary (Fig. 3E-3G in main text; table S4). We used lower AIC values as indication of a better fit model, and estimated Akaike weights ($w$) for each model to estimate the relative likelihood of the model, where models with higher $w$ have more support, and models with similar $w$ values are equally supported by the data [75].

### g. SENSITIVITY ANALYSES

We note that while niche and competitive differences are compound parameters derived from invading and monoculture growth rates measured for each lineage [25,28], we could not propagate uncertainty from individual growth rate estimates as the experiment was not designed to capture uncertainty within any genotype pair. This was a necessary trade-off in our experimental design, in which replication was limited within each genotype but was maximized across many genotype pairs to capture uncertainty in the relationship between ecological differences and genetic distance. We conducted several sensitivity analyses to assess the robustness of models testing how niche differences, competitive differences, and the potential for coexistence varied with the pairwise genetic distance of *S. polyrhiza* lineages.

First, we visually detected two, extreme outliers in our estimates of niche difference and coexistence among *S. polyrhiza* lineages. We removed these outliers prior to subsequent analyses, but the results remained qualitatively unchanged (fig. S12; table S8).

Second, the pairwise and phylogenetically structured nature of the dataset means that estimates of niche differences, competitive differences, and coexistence are non-independent. To account for the covariance in these traits between pairs of lineages, we ran a modified phylogenetic generalized least squares (PGLS) model in a Bayesian approach using the *phylopairs* package [36]. Here, covariance between lineages in a pairwise trait (e.g., competitive difference) is modelled explicitly using a covariance matrix based on ultrametric trees, with the assumption that some underlying trait (e.g., a resource acquisition trait) with phylogenetic signal is generating this pairwise covariance. We ran PGLS models using the *linreg.stan* function with standardized (z-scored) pairwise genetic distance as a predictor and temporal block as a covariate. We used default, weakly informative priors including Gaussian priors for predictor variables ($\mu = 0$; $\sigma = 10$), lognormal priors for the scale of the phylogenetic component of residual covariance ($\mu = -1$; $\sigma = 1$), and cauchy priors for the independent component of residual covariance ($\mu = 0$; $\sigma = 2.5$). We ran all models for 6000 iterations across four chains. We note that we could not additionally include experimental group IDs due to model complexity in these analyses, although we found from corresponding GAMMs that the variance explained by the experimental group was small (ranging from 2.3% to 4.3% of the total variance). We report the mean posterior estimate and the 95% Highest Posterior Density Intervals (HPDIs), the latter of which is defined as the narrowest interval that contains 95% of the probability mass (fig. S4 and S5).

Third, we tested if pairwise climate dissimilarity among experimental lineages could be a possible driver of divergent selection for niche and competitive differences, and therefore the potential for coexistence. To do this, we first extracted the mean and variability (SD) in annual lake surface water temperatures (LSWT) at each sampling locality from a published dataset based on satellite-derived temperatures [76]. LSWT has been previously used to accurately predict the geographical distribution and patterns of co-occurrence of duckweed species, including for *S. polyrhiza* [62,77]. Using the mean and SD in annual LSWT, we derived Gaussian curves to describe the climate niche of each lineage. We then estimated climatic niche overlap $\cap_C$ between pairs of lineages as the intersection of their Gaussian curves, where $\cap_C$ ranges 0 (no climatic niche overlap) to 1 (complete climatic niche overlap). Finally, we estimated pairwise climate dissimilarity between lineages as $1 - \cap_C$. We note that LWST data was not available for one lineage of *S. polyrhiza*. Estimates of climate dissimilarity therefore could not be obtained for competing pairs involving this lineage and thus were removed from climate analyses.

As climate dissimilarity was positively correlated with pairwise genetic distance (Pearson's $r$ = 0.240, $p < 0.001$; fig. S1B), we fit a linear effect of climate dissimilarity with pairwise genetic distance as a covariate using GAMMs. We included temporal block and experimental group as random effects, and fit separate models for niche differences, competitive differences, and coexistence as described previously. While we found a weakly positive and marginally significant effect of climate dissimilarity on niche differences ($\beta = 0.042$, 95% CI: -0.003 to 0.087, $p = 0.064$; fig. S7; table S5), we did not find a significant association between climate dissimilarity and competitive differences ($\beta = 0.154$, 95% CI: -0.062 to 0.370, $p = 0.153$; fig. S7; table S5) or the potential for coexistence among diverging populations ($\beta = 0.046$, 95% CI: -0.016 to 0.108, $p = 0.134$; fig. S7; table S5). We note, however, that the effect of genetic distance on niche differences and potential for coexistence remained significant even after accounting for climate dissimilarity as a covariate (table S5).

### SUPPLEMENTAL FIGURES

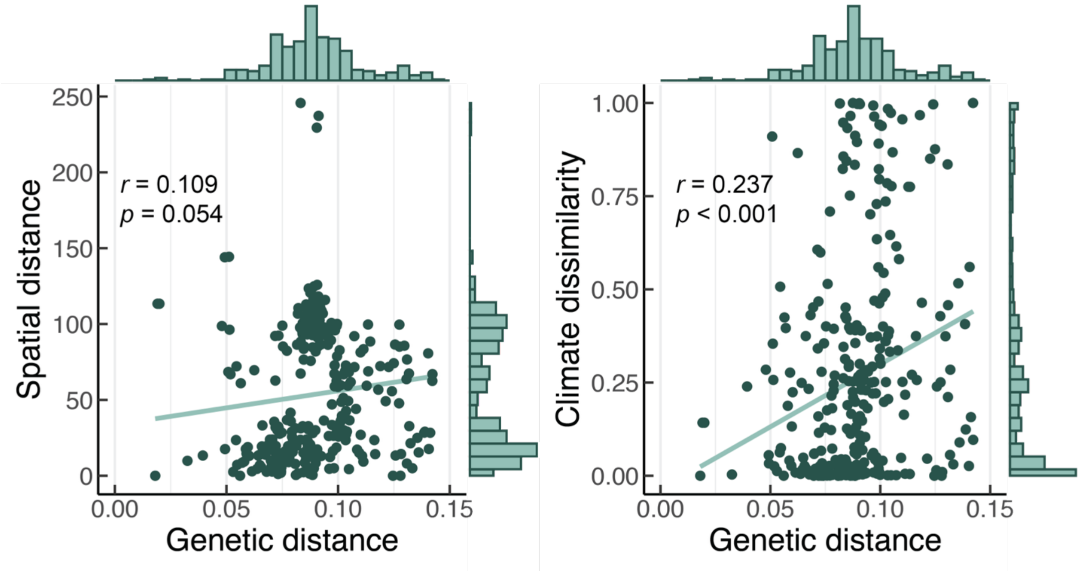


**Fig. S1.**

**Correlation between pairwise genetic distance and spatial or climate distance among *S. polyrhiza* lineages chosen for experimentation.** Data points and histograms represent estimates of genetic distance and: **(A)** spatial distance; or **(B)** climate distance (i.e., 'climate dissimilarity') for each competing pair of *S. polyrhiza*. The green line represents the Pearson's correlation coefficient ($r$) in each panel ($r_{spatial} = 0.109$, $p = 0.054$; $r_{climate} = 0.237$, $p < 0.001$). Note that genetic distance represents the proportion of nucleotide differences between lineages (substitutions per site). Spatial distance represents the Euclidean distance in spatial coordinates between each pair of allopatric lineages. Climate dissimilarity between pairs of lineages is 1 – the climatic niche overlap based on annual lake surface water temperatures (see **g. Sensitivity Analyses** for details).

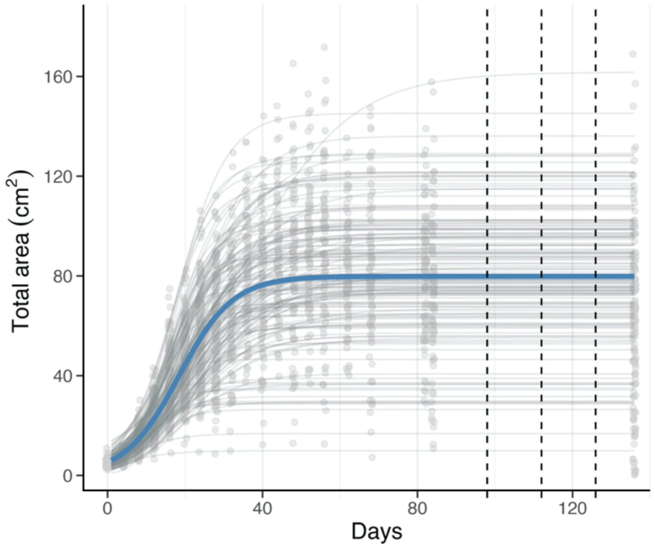


**Fig. S2.**

**Logistic growth fit for resident *Spirodela* lineages.** Population density was estimated as frond area ($cm^2$). Frond area was quantified by photographing a sub-sample of each resident lineage and calculating the surface area of living (green) fronds. We estimated population density for each resident lineage every 2-4 days before the competition trials (data points). We also estimated population density on the final day of the last competition trial. The logistic model fit across all resident lineages is given by the dark blue line ($K$ = 79.86 $cm^2$; $r$ = 0.143/day). Logistic model fits for each resident lineage are given by the thin grey lines. Vertical dashed lines represent the time point at which competition trials began in the three temporal blocks.

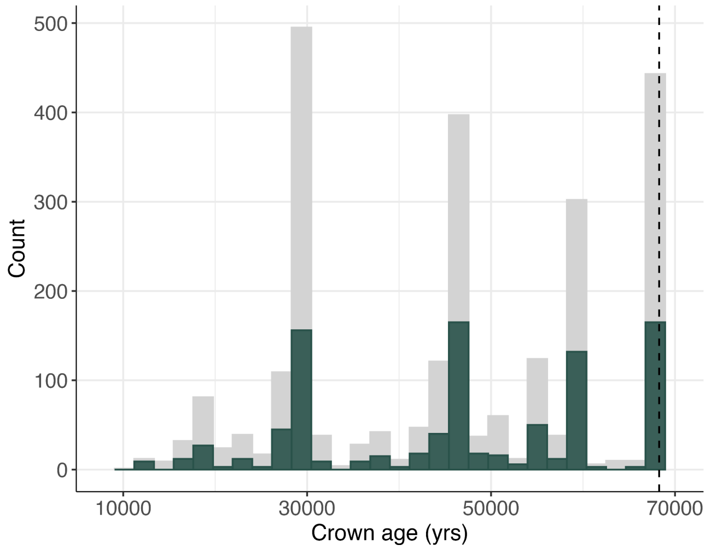


**Fig S3.**

**Distribution of divergence time (yrs) among *S. polyrhiza* lineages.** Gray bars show the distribution of divergence times for all possible pairwise combinations of lineages, while green bars show the distribution of divergence times for pairs of lineages for which we experimentally competed to test coexistence. The dashed vertical line shows the maximum divergence time (68,000 years) estimated among *S. polyrhiza* lineages in our experiment.

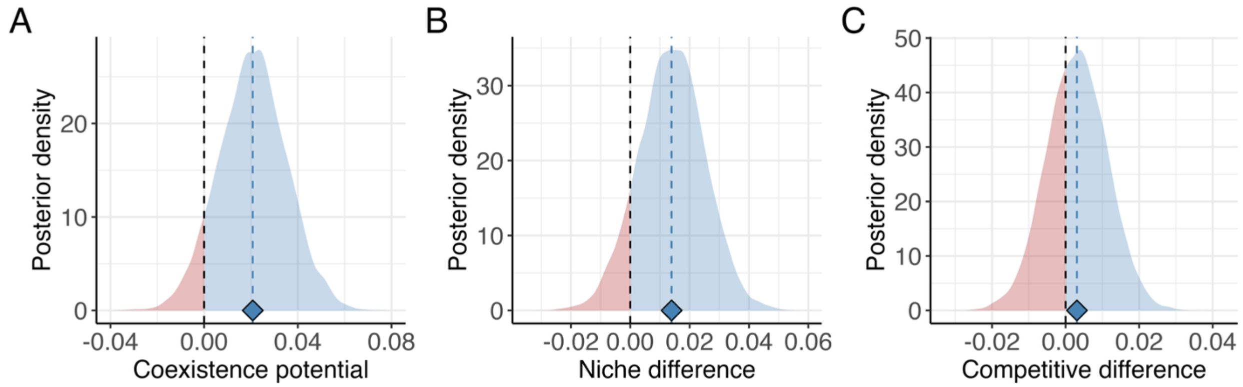


**Fig. S4**.

**Posterior distribution of slope estimates from models accounting for phylogenetic non-independence.** Panels show the posterior distribution of the slope (across pairwise genetic distance) for: **(A)** coexistence potential; **(B)** niche differences; or **(C)** competitive differences in pairs of *S. polyrhiza* lineages. Diamonds and dashed blue lines represent the mean posterior estimate of the slope. Black dashed lines indicate the value at which the posterior slope is zero. Negative and positive posterior values are in red and blue, respectively.

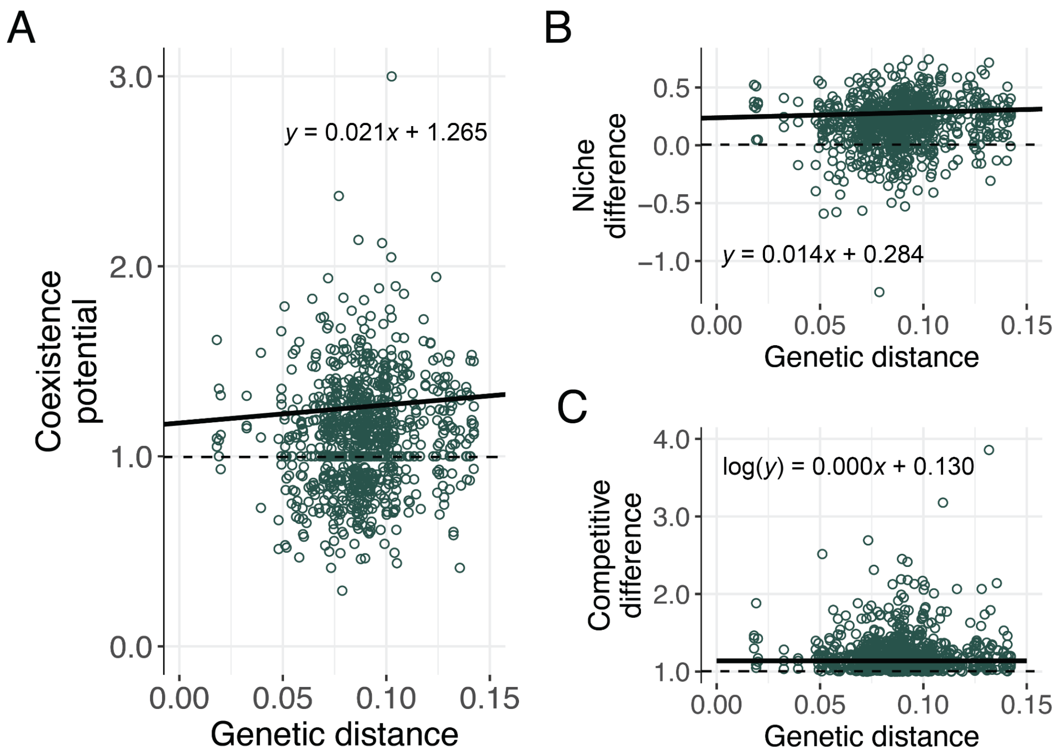


**Fig. S5.**
**Accounting for phylogenetic non-independence when testing for changes in niche differences, competitive differences, and coexistence potential with genetic distance.** Panels show the relationship between pairwise genetic distance and: **(A)** coexistence potential; **(B)** niche differences; or **(C)** competitive differences across replicate pairs of competing *S. polyrhiza* lineages. Slope estimates (dark line) represent the mean posterior estimate of the slope obtained from a Bayesian, modified phylogenetic generalized least squares (PGLS) model. Competitive differences were log transformed. Note that genetic distance represents the proportion of nucleotide differences between lineages (substitutions per site).

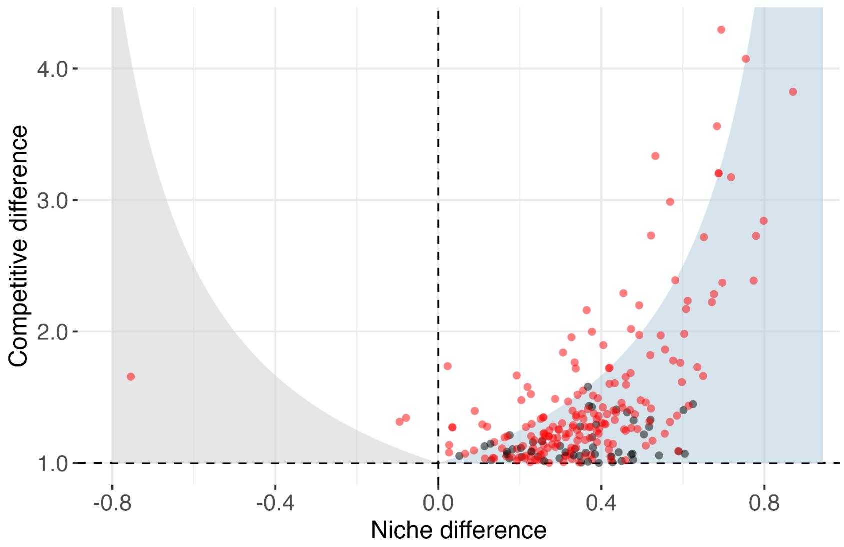


**Fig. S6.**

**Competitive dominance of *S. intermedia* versus *S. polyrhiza*.** Data points represent pairwise estimates of niche and competitive differences plotted relative to the zone of potential coexistence (blue shading). Data points are plotted at the replicate level and are coloured by whether *S. intermedia* are competitively dominant over *S. polyrhiza* (red), or if *S. polyrhiza* are competitively dominant over *S. intermedia* (black).

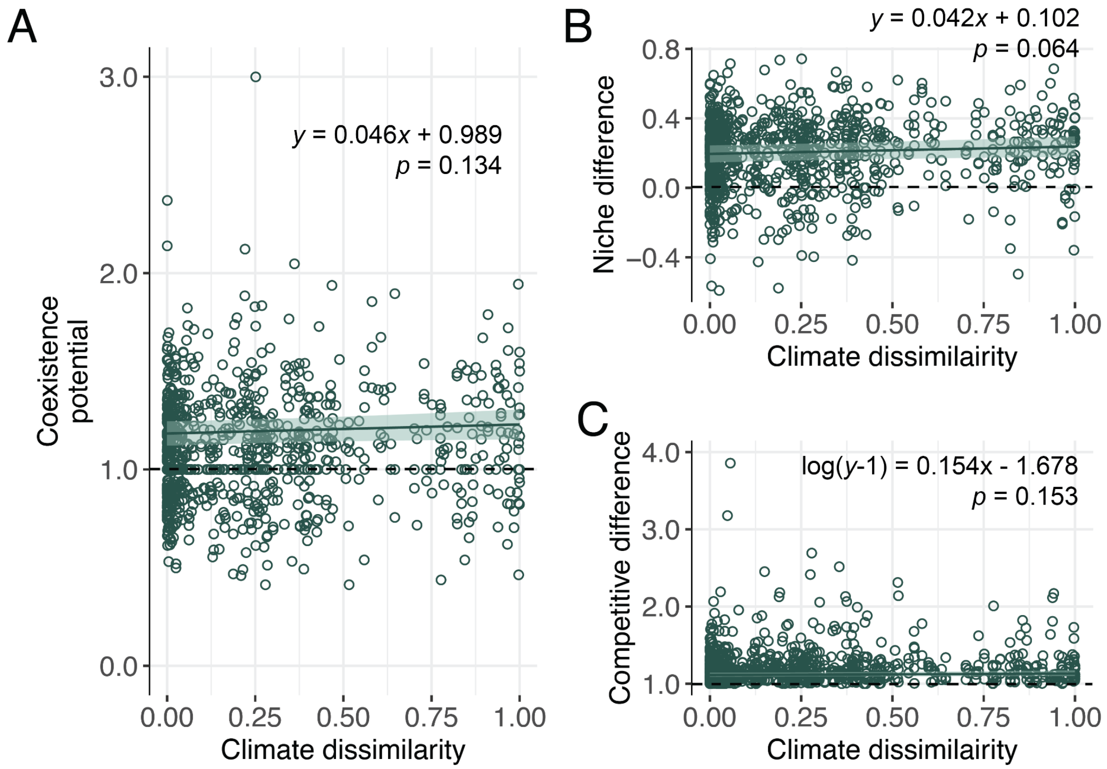


**Fig. S7.**
**Changes in niche differences, competitive differences, and coexistence potential with climate dissimilarity.** Panels show the relationship between pairwise climate dissimilarity and (**A**) coexistence potential; **(B)** niche differences; or (**C**) competitive differences for replicate pairs of competing *S. polyrhiza* lineages. Note that competitive differences use a shifted response (*y*-1) and are modelled using a Tweedie distribution with a log link. Model predicted slopes (dark green line), 95% confidence intervals (light green bands), and *p* values for the slope term are shown (see table S5 for model coefficients). Climate dissimilarity between pairs of lineages is 1 – the climatic niche overlap based on annual lake surface water temperatures (see **g. Sensitivity Analyses** for details).

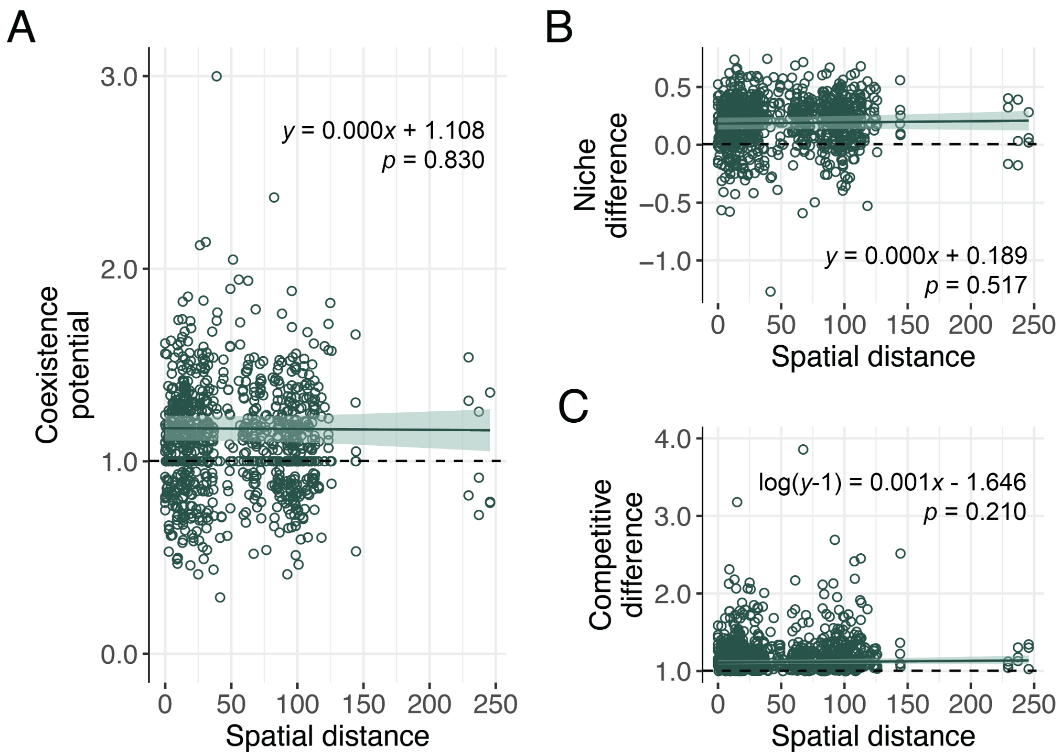


**Fig. S8.**

**Changes in niche differences, competitive differences, and coexistence potential with spatial distance.** Panels show the relationship between pairwise spatial distances and **(A)** coexistence potential; **(B)** niche differences; or **(C)** competitive differences for replicate pairs of competing *S. polyrhiza* lineages. Note that competitive differences use a shifted response (*y*-1) and are modelled using a Tweedie distribution with a log link. Model predicted slopes (dark green line), 95% confidence intervals (light green bands), and *p* values for the slope term are shown (see table S1 for model coefficients). Spatial distance represents the Euclidean distance in spatial coordinates between each pair of allopatric lineages.

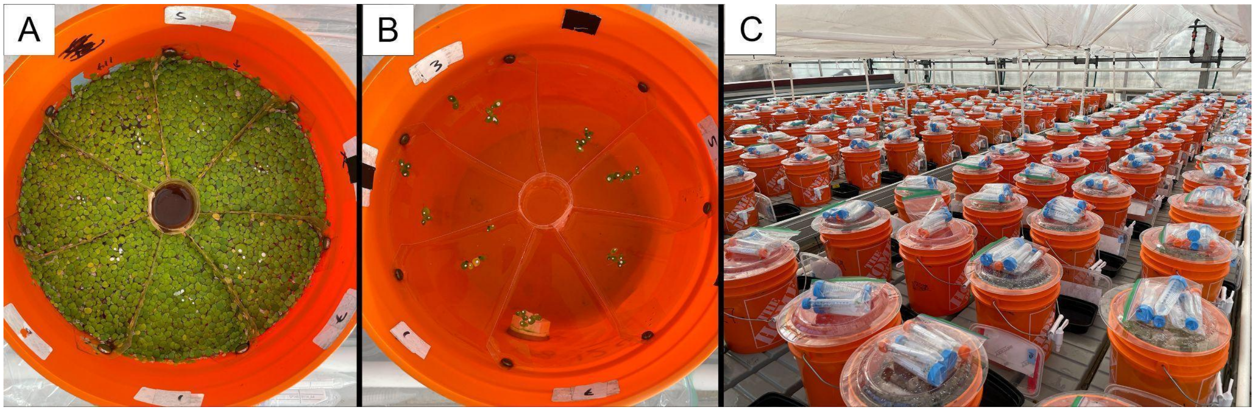


**Fig. S9.**

**Experimental setup for competition trials.** Panel **(A)** shows an example of a bucket where 'invader' lineages (white dots) are competing with a 'resident' lineage so that invasion growth rates ($g_{ij}$ and $g_{ji}$) can be estimated. Panel **(B)** shows an example of a bucket where each lineage is growing without competitors so that monoculture growth rates ($g_{i0}$) can be estimated. All buckets were installed with 5 cm deep plastic dividers at the surface to prevent crossover of genotypes. Panel **(C)** shows the full experimental set up for one temporal block.

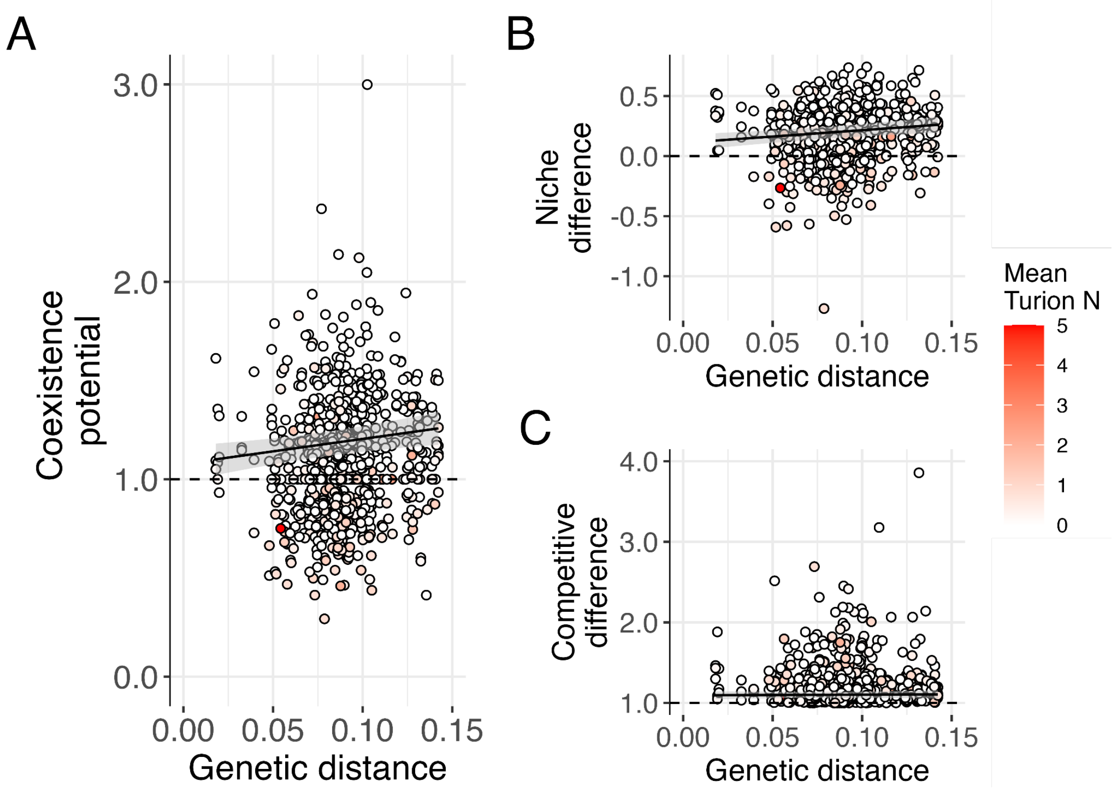


**Fig. S10.**

**Number of turions produced by each invading lineage of the competition trial.** Panels are the same as Figure 2 in the main text, except coloured by the average number of turions produced over 8 days for each invading individual during the competition trial. Most invaders produced no turions (white circles). If produced, turion number does not show any biases across genetic distance, and does not contribute to large values in (A) coexistence potential, (B) niche difference, or (C) competitive difference.

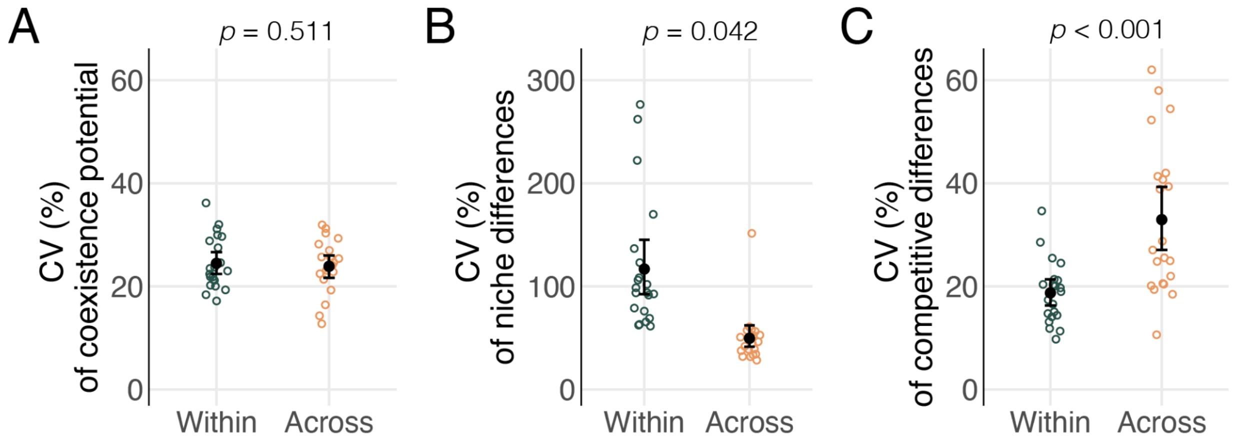


**Fig. S11.**
**Comparing variability in niche differences, competitive differences, and coexistence potential within and between the species boundary.** Variability was measured as the coefficient of variation (CV; %) for: **(A)** coexistence potential; **(B)** niche differences; or **(C)** competitive differences. Green and orange circles show estimates of variability within versus across the species boundary (i.e., variability among *S. polyrhiza* pairs or among *S. polyrhiza*-*S. intermedia* pairs, respectively). Error bars represent 95% CIs obtained from bootstrap sampling. Note that each point represents an estimate of CV for an experimental group (with each group comprising six competing and unique lineages). *P* values are shown for tests of differences in CV within versus across the species boundary (table S7; see Supplementary Methods for details).

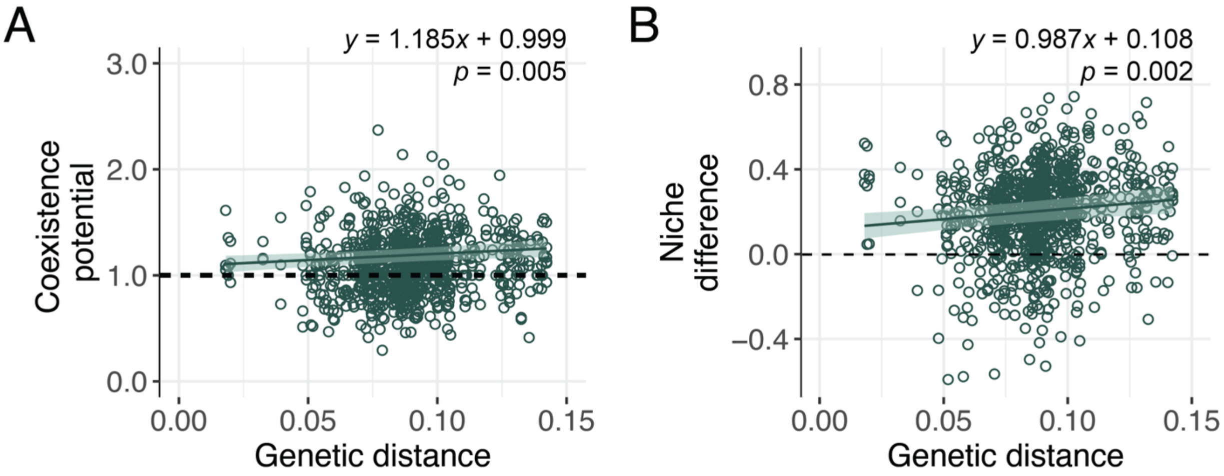


**Fig. S12**.

**Removing outliers in coexistence potential and niche differences.** Panels show the relationship between pairwise genetic distance and: **(A)** coexistence potential; or **(B)** niche differences for replicate pairs of competing *S. polyrhiza* lineages. Sensitivity tests of removing outliers were not conducted for competitive differences as outliers were not visually detected. Model predicted slopes (dark green line), 95% CIs (green bands), and *p* values for the slope term are shown (see table S8 for model coefficients). Note that genetic distance represents the proportion of nucleotide differences between lineages (substitutions per site).

### SUPPLEMENTAL TABLES

**Table S1.**

**Analyses of niche differences, competitive differences, and coexistence potential with spatial distance among *S. polyrhiza* lineages.** Separate GAMMs were fit for each response variable with a linear effect of pairwise spatial distance, and with experimental group and temporal block as random effects. Spatial distance represents the Euclidean distance in spatial coordinates between each pair of allopatric lineages. LCI and UCI stand for lower and upper 95% CIs. Note that coefficients for competitive differences are given on the log link scale. *Italicised* are significant terms where $p < 0.05$.

| Response | Term | Estimate | SE | *t*-value | *p* | LCI | UCI |
|---|---|---|---|---|---|---|---|
| Niche differences | *Intercept* | *0.189* | *0.041* | *4.555* | *<0.001* | *0.097* | *0.280* |
| | Slope | 0.000 | 0.000 | 0.648 | 0.517 | 0.000 | 0.000 |
| Competitive difference | *Intercept* | *-1.646* | *0.226* | *-7.275* | *<0.001* | *-2.147* | *-1.145* |
| | Slope | 0.001 | 0.001 | 1.254 | 0.210 | -0.001 | 0.002 |
| Coexistence potential | *Intercept* | *1.108* | *0.071* | *15.623* | *<0.001* | *0.951* | *1.266* |
| | Slope | 0.000 | 0.000 | -0.215 | 0.830 | 0.000 | 0.000 |

**Table S2.**

**Analyses of niche differences, competitive differences, and coexistence potential across genetically diverging *S. polyrhiza* lineages.** Separate GAMMs were fit to each response variable with a linear effect of pairwise genetic distance (nucleotide differences between lineages), and with experimental group and temporal block as random effects. LCI and UCI stand for lower and upper 95% CIs. Note that coefficients for competitive differences are given on the log link scale. *Italicised* are significant terms where $p < 0.05$.

| Response | Term | Estimate | SE | *t*-value | *p* | LCI | UCI |
|---|---|---|---|---|---|---|---|
| Niche difference | *Intercept* | *0.100* | *0.050* | *2.008* | *0.045* | *-0.007* | *0.207* |
| | *Slope* | *1.058* | *0.330* | *3.206* | *0.001* | *0.397* | *1.718* |
| Competitive difference | *Intercept* | *-1.672* | *0.260* | *-6.433* | *<0.001* | *-2.235* | *-1.109* |
| | Slope | 0.835 | 1.512 | 0.552 | 0.581 | -2.196 | 3.867 |
| Coexistence potential | *Intercept* | *0.994* | *0.080* | *12.466* | *<0.001* | *0.821* | *1.167* |
| | *Slope* | *1.262* | *0.433* | *2.917* | *0.004* | *0.394* | *2.130* |

**Table S3.**

**Analyses of whether mean niche differences, competitive differences, and coexistence potential differ within (i.e., *S. polyrhiza*–*S. polyrhiza* pairs) versus across the species boundary (i.e., *S. polyrhiza*–*S. intermedia* pairs).** Separate GAMMs were fit for each response variable with a linear effect of species level (within or across species), and with experimental group and temporal block as random effects. LCI and UCI stand for lower and upper 95% CIs. Intercepts represent ‘within’ species estimates. Note that coefficients for competitive differences are given on the log link scale. *Italicised* are significant terms where $p < 0.05$.

| Response | Term | Estimate | SE | *t*-value | *p* | LCI | UCI |
|---|---|---|---|---|---|---|---|
| Niche differences | *Intercept* | *0.194* | *0.029* | *6.706* | *<0.001* | *0.130* | *0.258* |
| | *Across* | *0.149* | *0.015* | *10.240* | *<0.001* | *0.120* | *0.179* |
| Competitive difference | *Intercept* | *-1.599* | *0.260* | *-6.145* | *<0.001* | *-2.177* | *-1.022* |
| | *Across* | *0.641* | *0.066* | *9.658* | *<0.001* | *0.508* | *0.774* |
| Coexistence potential | *Intercept* | *1.106* | *0.063* | *17.491* | *<0.001* | *0.966* | *1.247* |
| | *Across* | *0.106* | *0.020* | *5.222* | *<0.001* | *0.065* | *0.147* |

**Table S4.**

**Model comparison for linear versus non-linear models of niche differences, competitive differences, and coexistence potential across the species boundary (i.e., across *S. polyrhiza-S. intermedia* pairs).** Separate GAMMs were fit for each response variable across pairwise genetic distance (nucleotide differences between lineages). Non-linear effects were fit using a spline with a restricted basis dimension ($k = 3$). For each model, AIC values, differences in AIC (ΔAIC) from the best model (*italicised*), and Akaike weights (*w*) are reported.

| Response | Model type | AIC | ΔAIC | Akaike *w* |
|---|---|---|---|---|
| Niche difference | Linear | -510.735 | 2.283 | 0.242 |
| | *Non-linear* | *-513.018* | *0.000* | *0.758* |
| Competitive difference | *Linear* | *-899.789* | *0.000* | *0.502* |
| | Non-linear | -899.776 | 0.013 | 0.498 |
| Coexistence potential | Linear | 258.691 | 4.085 | 0.115 |
| | *Non-linear* | *254.606* | *0.000* | *0.885* |

**Table S5.**

**Analyses of niche differences, competitive differences, and coexistence potential with pairwise climate dissimilarity among *S. polyrhiza* lineages.** Separate GAMMs were fit for each response variable with a linear effect of pairwise climate dissimilarity (1 – climatic niche overlap) and pairwise genetic distance (nucleotide differences between lineages). Experimental group and temporal block were fit as random effects. LCI and UCI stand for lower and upper 95% CIs. Note that coefficients for competitive differences are given on the log link scale. *Italicised* are significant terms where $p < 0.05$.

| Response | Term | Estimate | SE | *t*-value | *p* | LCI | UCI |
|---|---|---|---|---|---|---|---|
| Niche difference | *Intercept* | *0.102* | *0.048* | *2.118* | *0.034* | *-0.002* | *0.206* |
| | Climate | 0.042 | 0.023 | 1.856 | 0.064 | -0.003 | 0.087 |
| | *Genetic* | *0.940* | *0.326* | *2.886* | *0.004* | *0.287* | *1.592* |
| Competitive difference | *Intercept* | *-1.678* | *0.259* | *-6.477* | *<0.001* | *-2.239* | *-1.117* |
| | Climate | 0.154 | 0.107 | 1.431 | 0.153 | -0.062 | 0.370 |
| | Genetic | 0.413 | 1.551 | 0.266 | 0.790 | -2.694 | 3.521 |
| Coexistence potential | *Intercept* | *0.989* | *0.079* | *12.539* | *<0.001* | *0.817* | *1.160* |
| | Climate | 0.046 | 0.031 | 1.499 | 0.134 | -0.016 | 0.108 |
| | *Genetic* | *1.207* | *0.440* | *2.741* | *0.006* | *0.324* | *2.090* |

**Table S6.**

**Nutrient composition for freshwater medium used for competition trials.** Nutrient media originally formulated by Docauer [58]. This media mimics the nutrient contents, pH, and conductivity of freshwater ponds and lakes containing duckweeds including *Spirodela polyrhiza*.

| Chemical | Stock concentration | Stock per L media | Final media concentration | |
|---|---|---|---|---|
| | (g/L) | (mL) | (mg/L) | (mM) |
| $Na_2EDTA.2H_2O$ | 20.00 | 2.0 | | 0.107 |
| $NaNO_3$ | 25.00 | 2.0 | 8.2 N | 0.588 |
| $K_2HPO_4$ | 3.68 | 1.0 | 0.7 P | 0.021 |
| KCl | 50.00 | 1.0 | 26.2 K | 0.671 |
| $CaCl_2$ | 25.00 | 2.0 | 18.1 Ca | 0.451 |
| $MgSO_4.7H_2O$ | 37.50 | 0.5 | 1.8 Mg | 0.076 |
| Micronutrients | See below | 1.0 | See below | |
| $NaHCO_3$ | 60.00 | 1.0 | 43.6 $HCO_3$ | 0.714 |

| Micronutrient Chemical | Stock concentration | Final media concentration | |
|---|---|---|---|
| | (g/100mL) | (mg/L) | (mM) |
| $FeSO_4.7H_2O$ | 0.995 | 2.0 Fe | 0.036 |
| $MnCl_2.4H_2O$ | 0.072 | 0.2 Mn | 0.004 |
| $Na_2MoO_4.2H_2O$ | 0.044 | 0.2 Mo | 0.002 |
| $H_3BO_3$ | 0.057 | 0.1 B | 0.009 |
| $ZnSO_4.7H_2O$ | 0.044 | 0.1 Zn | 0.002 |
| $Na_2EDTA.2H_2O$ | 2.000 | | 0.054 |
| *Mix the following separately and add 1 mL per 100 mL of micronutrient stock:* | | | |
| $CuSO_4.5H_2O$ | 0.004 | 0.0001 Cu | 0.0000016 |
| $CoCl_2.6H_2O$ | 0.400 | 0.01 Co | 0.00017 |
| $Na_2EDTA.2H_2O$ | 0.400 | | 0.00011 |

**Table S7.**

**Analysis of whether variability in niche differences, competitive differences, and coexistence potential differ within (i.e., *S. polyrhiza*–*S. polyrhiza* pairs) versus across the species boundary (i.e., *S. polyrhiza*–*S. intermedia* pairs).** Separate Modified signed-likelihood ratio tests (MLRTs) were fit for each response variable with species level ('within' versus 'across') as a predictor to test for equality in the coefficient of variation (CV). *Italicised* are significant terms where $p < 0.05$.

| Response | MLRT | *p* |
|---|---|---|
| *Niche differences* | *4.128* | *0.042* |
| *Competitive differences* | *16.07* | *<0.001* |
| Coexistence potential | 0.424 | 0.515 |

**Table S8.**

**Removal of outliers when testing for changes in niche differences and coexistence potential across genetically diverging *S. polyrhiza* lineages.** Separate GAMMs were fit for each response variable with a linear effect of pairwise genetic distance (nucleotide differences between lineages), and with experimental group and temporal block as random effects. LCI and UCI stand for lower and upper 95% CIs. *Italicised* are significant terms where $p < 0.05$. Note that outliers were not detected for estimates of competitive differences.

| Response | Term | Estimate | SE | *t*-value | *p* | LCI | UCI |
|---|---|---|---|---|---|---|---|
| Niche difference | *Intercept* | *0.108* | *0.049* | *2.219* | *0.027* | *0.003* | *0.213* |
| | *Slope* | *0.987* | *0.321* | *3.077* | *0.002* | *0.343* | *1.630* |
| Coexistence potential | *Intercept* | *0.999* | *0.080* | *12.477* | *<0.001* | *0.825* | *1.173* |
| | *Slope* | *1.185* | *0.420* | *2.822* | *0.005* | *0.343* | *2.028* |
